# Dissecting the Functional Organization of the *C. elegans* Serotonergic System at Whole-Brain Scale

**DOI:** 10.1101/2023.01.15.524132

**Authors:** Ugur Dag, Ijeoma Nwabudike, Di Kang, Matthew A. Gomes, Jungsoo Kim, Adam A. Atanas, Eric Bueno, Cassi Estrem, Sarah Pugliese, Ziyu Wang, Emma Towlson, Steven W. Flavell

**Author notes:** These authors contributed equally to this work (Ugur Dag, Ijeoma Nwabudike, Di Kang).

## Abstract

Serotonin controls many aspects of animal behavior and cognition. But how serotonin acts on its diverse receptor types in neurons across the brain to modulate global activity and behavior is unknown. Here, we examine how serotonin release from a feeding-responsive neuron in *C. elegans* alters brain-wide activity to induce foraging behaviors, like slow locomotion and increased feeding. A comprehensive genetic analysis identifies three core serotonin receptors that collectively induce slow locomotion upon serotonin release and three others that interact with them to further modulate this behavior. The core receptors have different functional roles: some induce behavioral responses to sudden increases in serotonin release, whereas others induce responses to persistent release. Whole-brain calcium imaging reveals widespread serotonin-associated brain dynamics, impacting different behavioral networks in different ways. We map out all sites of serotonin receptor expression in the connectome, which, together with synaptic connectivity, helps predict serotonin-associated brain-wide activity changes. These results provide a global view of how serotonin acts at defined sites across a connectome to modulate brain-wide activity and behavior.

## INTRODUCTION

Serotonin signaling is evolutionarily ancient and is critical for the control of behavior and cognition. In humans, dysfunction of the serotonergic system has been implicated in major depressive disorder, anxiety disorders, and other psychiatric diseases. It is the most common target of psychiatric drugs, and many psychotropic chemicals like psilocybin and LSD exert their effects via action on serotonin receptors. Efforts to understand how serotonin signaling controls behavior and cognition have been hindered by its complexity. The neurons that release serotonin are functionally diverse, they project broadly throughout the brain, and they exert their effects via 14 different receptors that have distinct functions and complex interactions. Developing an integrative framework for serotonergic function that relates anatomy, receptors, circuits, and behavior at the scale of a whole nervous system would greatly aid our understanding of this important neuromodulatory system.

In the mammalian brain, serotonin is released by many molecularly distinct subtypes of serotonergic neurons in the Raphe nuclei, each with their own widespread projections to distal brain regions (Okaty et al., 2020, 2019; Ren et al., 2019, 2018). Neural recordings from this heterogeneous population have revealed diverse activity profiles, reflecting rewarding stimuli like food, as well as complex cognitive processes (Cohen et al., 2015; Grossman et al., 2022; Li et al., 2016; Marques et al., 2020; Paquelet et al., 2022). This variety likely reflects the diverse inputs onto these neurons (Weissbourd et al., 2014). The outputs of the serotonergic neurons are varied and complex as well: optogenetic stimulation of serotonergic neurons in rodents can elicit many behavioral effects, including slow locomotion, waiting/perseverance, and changes in reward learning (Cazettes et al., 2021; Grossman et al., 2022; Lottem et al., 2018; Ren et al., 2018; Seo et al., 2019). Pharmacological and genetic studies have shown that serotonin acts through many distinct receptors to control different aspects of behavior (Donaldson et al., 2013; Yohn et al., 2017). Efforts to map out sites of serotonin receptor expression in circuits that receive dense serotonergic afferents show that many distinct cell types in a microcircuit can express distinct serotonin receptors (Bocchio et al., 2016; Puig and Gulledge, 2011). Slice recordings of individual cell types paired with pharmacology have shown bulk effects of serotonin application on individual cell types (Béïque et al., 2007; Goodfellow et al., 2009). However, these studies do not necessarily predict in vivo changes in circuit activity that accompany serotonin release (Puig et al., 2010). In the face of these complexities, it has been challenging to develop a mechanistic understanding of how dynamic serotonin release in live animals engages specific serotonin receptors to alter ongoing neural dynamics and behavior. Such a level of understanding will be critical in order to optimally target serotonin receptors to treat psychiatric disease.

It should be feasible to address these complexities in the nematode *C. elegans*, whose nervous system consists of 302 uniquely identifiable neurons with known connectivity (Cook et al., 2019; White et al., 1986; Witvliet et al., 2021). The serotonergic system plays an important role in controlling *C. elegans* behavior, and its molecular and anatomical organization resembles that of mammals. *C. elegans* utilizes the same synthesis pathway (*tph-1*/TPH, *bas-1*/AAAD), vesicle loading machinery (*cat-1*/VMAT), reuptake machinery (*mod-5*/SERT), and many of the same receptor types (5-HT1, 5-HT2, and more) found in mammals (Chase and Koelle, 2007). The few-to-many anatomical organization of the serotonergic system is also preserved in *C. elegans*: there are three pairs of serotonergic neurons (NSM, HSN, and ADF) and a much larger estimated number of neurons that express serotonin receptors (Chase and Koelle, 2007).

When *C. elegans* animals eat appetitive bacterial food, they display adaptive behavioral changes that are controlled by serotonin: slow locomotion, increased feeding, and increased egg-laying (Flavell et al., 2013; Horvitz et al., 1982; Iwanir et al., 2016; Sawin et al., 2000). While egg-laying is controlled by the serotonergic neuron HSN in a mostly isolated circuit in the *C. elegans* midbody, the other food-induced behavioral changes are controlled by the serotonergic neuron NSM in the head (Flavell et al., 2013; Iwanir et al., 2016; Rhoades et al., 2019; Sawin et al., 2000). NSM extends a sensory dendrite into the pharyngeal lumen and directly senses bacterial food that the animal is eating, such that NSM is only active while animals are eating bacteria (Rhoades et al., 2019). Its activity is strongly correlated with slow locomotion (Flavell et al., 2013; Iwanir et al., 2016; Rhoades et al., 2019) and the role of NSM becomes more prominent in fasted animals, an effect called the enhanced slowing response (Sawin et al., 2000). NSM appears to act through volume transmission: it releases serotonin at non-synaptic neurosecretory release sites that are adjacent to the nerve ring, the main neuropil of the *C. elegans* nervous system containing fibers from 196 neurons (Nelson and Colón-Ramos, 2013; White et al., 1986). Six *C. elegans* serotonin receptors have been identified so far, four of which have been suggested to influence locomotion and one of which influences feeding rates (Chase and Koelle, 2007; Morud et al., 2021). However, how these receptors are functionally organized to control behavior remains unknown. Given that these receptors are co-activated during serotonin volume transmission, how do they interact with one another? How is serotonin receptor expression arranged in the connectome and which sites of receptor expression drive which features of the behavioral change? And how does serotonin binding to this diverse set of receptors across distinct cell types alter the global brain dynamics that control behavior?

Here, we provide a global characterization of the serotonergic system that links these different scales of analysis. We find that serotonin release by NSM alters locomotion in a manner that depends on all six serotonin receptors. We decipher the functional roles of the receptors, and the interactions between them, by examining behavioral responses to serotonin release in a panel of 64 mutant strains that lack all possible combinations of receptors. This analysis also reveals that the receptors support widely different dynamical changes in behavior, with some receptors only supporting a transient behavioral change in response to increased serotonin release and others supporting a persistent behavioral change during continued serotonin release. Swapping the sites of receptor expression shows that the cell types expressing a receptor, rather than the molecular properties of the receptor, confer the kinetics of the behavioral response. We further map out the roles of receptor expression in individual cells, revealing that the kinetics of the behavioral response can be decomposed to individual cell types that support different temporal phases of the behavioral response to serotonin release. Finally, we layer these findings onto a brain-wide analysis of neural activity during serotonin release, which we map onto the physical wiring diagram of the animal. This reveals widespread NSM-associated neural dynamics across the brain that is reasonably well predicted by the serotonin receptor expression profiles and synaptic connectivity of the neurons. These results provide a global view of how serotonin acts on specific receptor types at defined sites across a connectome to modulate brain-wide activity and behavior.

## RESULTS

### An experimental platform to study serotonin-induced behavioral changes

To dissect the functional organization of the serotonergic system, we first developed an experimental platform where we could reliably induce a change in behavior that was completely dependent on serotonin signaling. The robust effect of NSM on locomotion during feeding provided a strong candidate system. When animals encounter and start eating bacterial food, NSM is activated and its activity correlates with slow locomotion (Iwanir et al., 2016; Rhoades et al., 2019). Previous studies have shown that NSM inactivation or serotonin depletion (via *tph-1* mutation) attenuates this feeding-induced behavior change, but it does not completely abolish it (Iwanir et al., 2016; Rhoades et al., 2019). Thus, while NSM serotonin strongly contributes to food-induced slowing, other pathways contribute as well. To obtain a system where serotonin acts on its own to alter neural activity and behavior, we optogenetically activated NSM with Chrimson in animals that were not consuming bacterial food. As expected, this induced a decrease in locomotion speed and head movements, as well as an increase in feeding (i.e. pharyngeal pumping), similar to when animals actually encounter food (Fig. 1A; defection and egg-laying behaviors were unaffected; see also Fig. S1A). These effects were abolished in *tph-1* mutants (Fig. 1A), suggesting that serotonin release is essential for this behavioral change.

**Figure 1.**
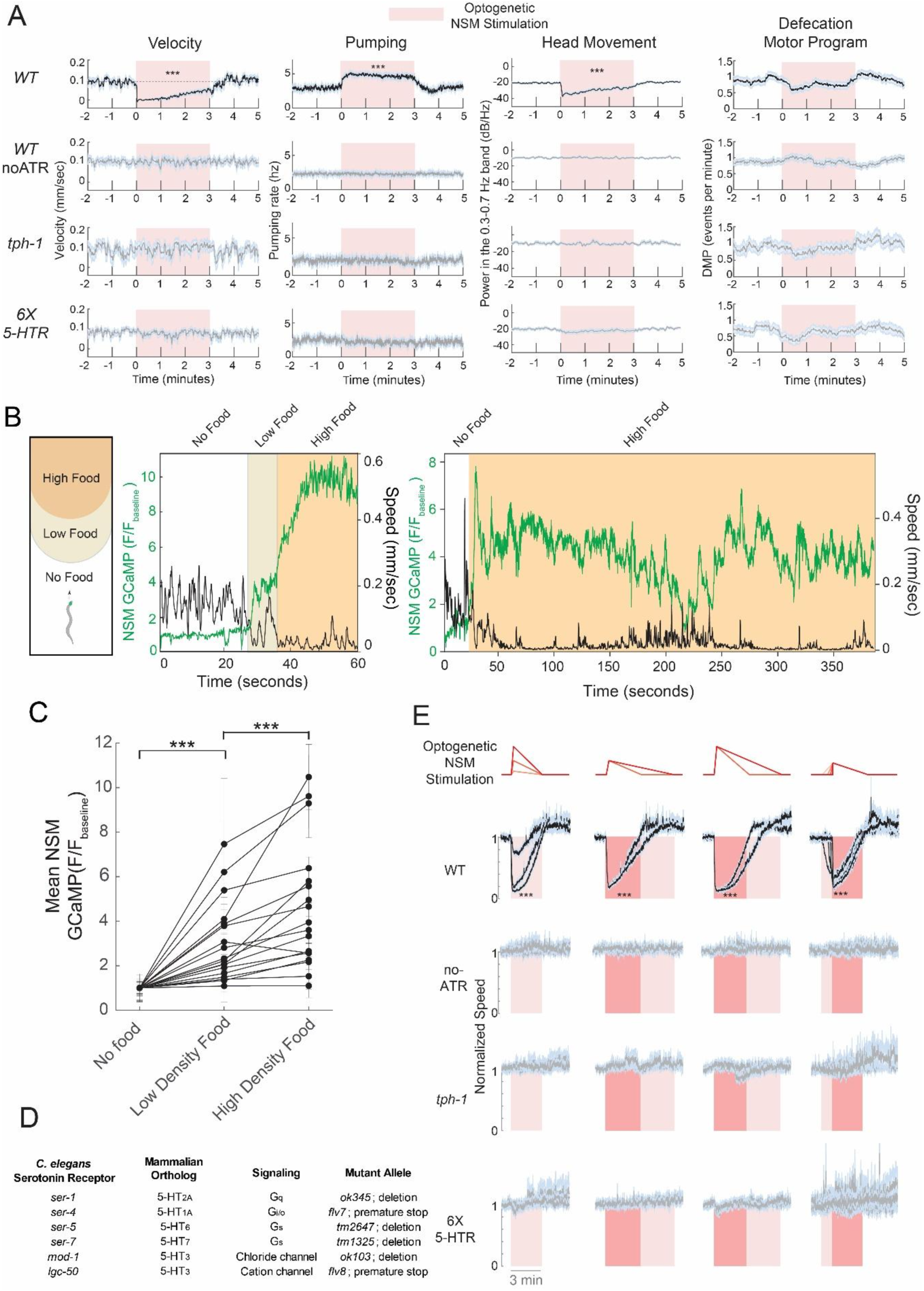
The serotonergic neuron NSM provides an experimental platform for studying serotonin-induced behavioral changes. (A) Behavioral changes evoked by NSM::Chrimson activation in WT, WT (no-ATR; lacking essential opsin co-factor, all-trans-retinal), *tph-1*, and 6x 5-HTR animals (animals with null mutations in all six *C. elegans* serotonin receptors). Experiments were performed on animals with no food on the plate (this is the case for all NSM optogenetics experiments in this study, unless otherwise stated). Four motor programs were quantified, as indicated: velocity, pharyngeal pumping (i.e. feeding), head movement (quantified as power in the frequency band corresponding to roaming speed, 0.3-0.7 Hz), and defecation motor programs (DMPs). NSM::Chrimson was activated in a pattern of a single block stimulus for three minutes. ***p<0.001, change in behavior during lights-on, Bonferroni-corrected t-test. N=24-78 stimulation trials per genotype (three per animal). Data shown as mean traces with error shading indicating standard error of the mean (SEM). (B) Left: cartoon depicting how animals navigate to bacterial food in this recording arena. Middle and Right: Two example traces showing simultaneous NSM GCaMP (green) and animal speed (black). Animals were recorded in an arena where they began the recording session in an area with no food, but they could navigate to a low-density food patch, inside of which was a high-density food patch. Shading indicates animal position in the arena. The example on the left shows NSM activity in each area of the arena (no-, low-, and high-density food). The example on the right (where the animal moved quickly into the high-density food region) shows an example of prolonged NSM activity in one area of the arena (high-density food). (C) Mean NSM GCaMP signal in each of three indicated regions of the recording arena. Data are shown as means ± standard deviation. Each line is one animal. ***p<0.001, paired t-test. n=16 animals. (D) A table of the six known *C. elegans* serotonin receptors and their properties. (E) Changes in animal speed induced by NSM::Chrimson activation with the indicated waveforms of light (top, red). Behavioral changes are shown for WT, WT (no-ATR), *tph-1*, and 6x 5-HTR. ***p<0.001, change in behavior during lights-on, Bonferroni-corrected t-test. N=104-518 animals per genotype. Data are shown as mean traces of normalized animal speed (animal speed, divided by pre-stimulus baseline speed) with error shading indicating SEM.

We considered which optogenetic stimulation patterns would most closely mimic natural patterns of NSM activity. Endogenous NSM activity during feeding consists of phasic bouts of activity that typically last for minutes. The amplitudes and durations of these activity bouts can vary (time range: from 1min to >10min; see Ji et al., 2021). To test whether the amplitudes of NSM activity bouts reflect important environmental cues, we recorded endogenous NSM activity in environments with varying external stimuli. We examined NSM GCaMP as animals encountered a bacterial food patch containing different densities of food arranged in concentric rings, with the densest food at the center (Fig. 1B-C). NSM activity increased in a manner that was time-locked to the initial food encounter. In addition, it increased further when animals subsequently encountered the denser part of the food patch. This indicates that the amplitudes of NSM activity bouts reflect food density. NSM activity bouts continue for many minutes (Fig. 1B, right), during which time they are anti-correlated with animal speed. Previous work has shown that NSM activity bouts eventually terminate in a manner that is closely aligned to the onset of roaming behavior (Ji et al., 2021). Thus, NSM activity is influenced by appetitive stimuli in the environment and is inversely correlated with animal speed.

Based on these results, we optogenetically activated NSM with different waveforms of light to mimic NSM activity bouts of different amplitudes and durations, directly parameterized to match our previous native recordings of NSM during food patch encounters (Rhoades et al., 2019). As above, these experiments (and all others below, unless otherwise indicated) were performed in the absence of food. Control experiments carried out with simultaneous NSM calcium imaging confirmed that these different optogenetic stimuli induced expected changes in NSM activity (Fig. S1B). Wild-type animals exhibited behavioral responses that closely tracked the NSM optogenetic stimulus: the magnitude of the reduction in speed scaled with optogenetic stimulus intensity, and the duration of the speed reduction scaled with the duration of the stimulus (Fig. 1E). Behavioral responses were abolished in serotonin-deficient *tph-1* mutant animals, indicating that serotonin is required for all these effects (Fig. 1E). In addition, these effects were also abolished in the sextuple mutant lacking all six known *C. elegans* serotonin receptors (*ser-1, ser-4, ser-5, ser-7, mod-1,* and *lgc-50*; Fig. 1D-E). This experimental paradigm allows us to evoke serotonin release with varying dynamics and examine how activation of a defined set of serotonin receptors modulates circuit activity to drive the behavioral response.

### Six *C. elegans* serotonin receptors interact to control locomotion

We next examined how each serotonin receptor contributes to the NSM-induced behavioral change. First, we crossed the NSM::Chrimson strain into the six single mutant strains that lack each of the serotonin receptors. The behavioral response of each mutant was attenuated compared to wild-type animals, though none of these mutants had fully abolished slowing (Fig. 2A-B; displayed as speed normalized to pre-stimulus baseline speeds; baseline speeds were all similar to wild-type, Fig. S2A). This indicates that each receptor plays some role in NSM-induced slowing. We next crossed the NSM::Chrimson strain into the six quintuple mutant strains that each has only a single serotonin receptor remaining (for simplicity, we refer to these strains as *mod-1*-only, *ser-1*-only, etc; though we note that there could be unidentified serotonin receptors in the *C. elegans* genome). Mutants with only *mod-1, ser-4,* or *lgc-50* intact all displayed NSM-induced slowing, though to varying degrees (Fig. 2A-B). Mutants with only *ser-5* and *ser-1* intact displayed almost no slowing, and mutants with only *ser-7* intact did not slow at all. Taken together, these data suggest that each of the six serotonin receptors contributes to NSM-induced slowing, though their roles likely differ. Comparing the single and quintuple mutants already suggested that there were likely to be complexities in how the serotonin receptors interact to control slowing. For example, the loss of *ser-1* led to a greater deficit in slowing than the loss of *ser-4*. However, *ser-4* on its own could promote robust slowing while *ser-1* could not (Fig. 2A-B). These results suggest that the serotonergic control of locomotion is mediated by activation of a set of six receptors that interact to modulate behavior.

**Figure 2.**
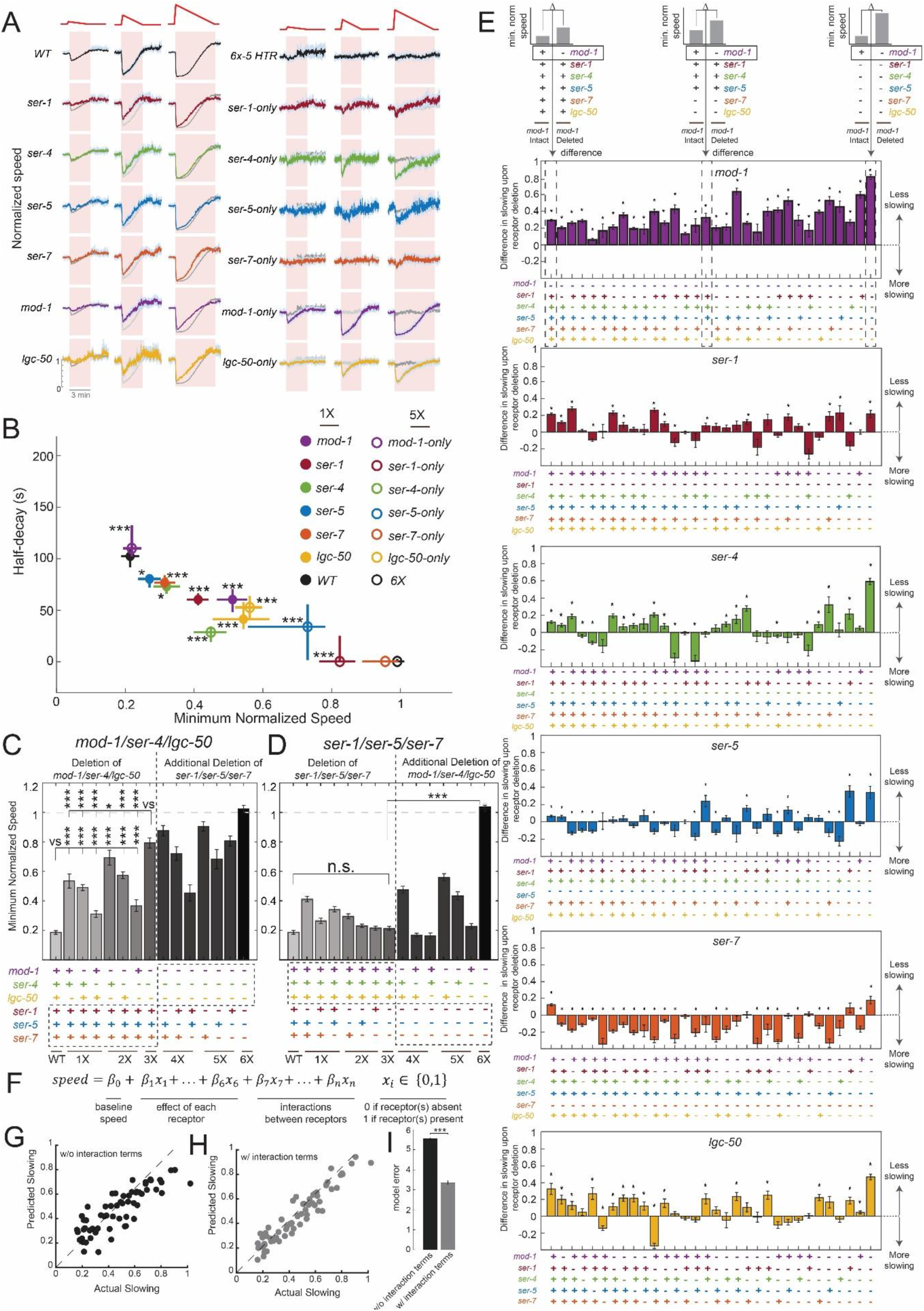
Comprehensive genetic analysis reveals how six serotonin receptors interact to drive the locomotion changes elicited by NSM activation. (A) Changes in animal speed induced by NSM::Chrimson activation with the indicated waveforms of light (top, red). Data are shown for WT, 6x 5-HTR (sextuple 5-HTR mutant), each serotonin receptor single mutant, and each quintuple mutant with a single serotonin receptor left intact. These results are quantified in panel (B). The gray traces in the lower rows show the results from the top row for reference (i.e. WT data for the left three columns and 6x 5-HTR for the right three columns). Data are shown as mean traces of normalized animal speed (animal speed, divided by pre-stimulus baseline speed) with error shading indicating SEM. Note that all of the summary analyses in this figure examine responses to the medium intensity stimulus. (B) Quantification of the behavioral traces in panel (A), for the medium intensity stimulus depicted in panel (A). x-axis shows the average slowing across animals (defined as minimum normalized speed during light stimulation for each animal); y-axis shows the decay rate of speed as it regresses back to baseline after the moment of maximum slowing (defined as time to decay back to 50% of baseline). For single mutants, *p<0.05, ***p<0.001 versus WT for both min speed and decay, Bonferroni-corrected empirical p-value that difference between distributions is non-zero. For quintuple mutants, ***p<0.001 versus 6x 5-HTR for min speed, Bonferroni-corrected empirical p-value that difference between distributions is non-zero. n=45-496 animals per genotype. Data are shown as means ± 95% confidence interval. (C) Minimum normalized speed during NSM stimulation for animals of the indicated genotypes, for the medium intensity stimulus depicted in panel (A). Colored dots in the x-axis labels indicate the presence or absence (i.e. mutation) of the indicated serotonin receptor: the presence of a dot indicates the presence of a receptor in the animal’s genotype. *p<0.05, ***p<0.001, empirical p-value that difference between distributions is non-zero. N=98-496 animals per genotype. (D) Minimum normalized speed during NSM activation for animals of the indicated genotypes, shown for the medium intensity stimulus. Results are displayed as in (C). ***p<0.001, empirical p-value that difference between distributions is non-zero. N=59-477 animals per genotypes. (E) Six plots depicting the change in NSM-induced slowing that is caused by a given receptor’s mutation, in the different genetic backgrounds (i.e. combinations of other serotonin receptor mutations) that were tested. These data are from the medium intensity optogenetic stimulus. The graphic at the top illustrates how the y-axis values of these plots were calculated: they reflect the difference in the level of slowing in the genotype indicated, compared to a corresponding genotype that is the same except the receptor in question is present. Data are shown as means ± SEM. Statistical test was a t-test for whether the receptor deletion altered slowing, compared to genetic background with receptor present. Asterisk indicates statistical significance at Benjamini-Hochberg FDR < 0.05 (corrected for all comparisons being performed for each receptor). Average n=278 animals per genotype. (F) Form of equation used to predict level of NSM-induced slowing of animals based on their serotonin receptor genotype. Interaction terms indicate the joint presence of two or more receptors (for them, 𝑥_!_ = 1 if all such receptors are present and 𝑥_!_ = 0 if one or more is absent). The interaction terms are only included in models as indicated, and their criteria for inclusion are described in the main text and Methods. The model was trained and tested on data from the medium intensity optogenetic stimulus, to avoid any issues of undetectable responses (with weak stimuli) or saturated responses (with strong stimuli). (G) Accuracy of the model shown in (F) with no interaction terms included. Accuracy is shown as the relationship between the actual mean level of slowing for a given genotype (x-axis) versus the model’s prediction of the level of slowing (y-axis). Each dot is a different genotype with different serotonin receptors present (64 total), and predicted slowing is from a 5-fold crossvalidation design, showing results from testing data. For crossvalidation, the allocation of animals into training and testing datasets was based on splitting the full pool of animals into five groups. (H) Model accuracy, shown as in (G). Here, the model is the one illustrated in (E), but with interaction terms included, as described in the text and Methods. (I) The model with interaction terms predicts slowing across the genotypes with higher accuracy. Data are the mean level of error ± standard deviations of five different models from a 5-fold crossvalidation design. ***p<0.001, t-test.

To decipher how these receptors interact, we crossed the NSM::Chrimson strain to the 64 different mutant backgrounds that lacked all possible combinations of the six serotonin receptors. We then quantified their locomotion changes in response to different patterns of NSM stimulation. We analyzed these datasets in several ways. First, we identified the subset of receptors that are required for slowing. The results across this full set of genotypes indicate that *mod-1, ser-4,* and *lgc-50* are the primary receptors that drive slow locomotion, so we refer to them as the “driver” receptors. The triple mutant lacking all three of these receptors displayed a slowing deficit that was almost as severe as the sextuple mutant, and the double mutants lacking combinations of these three receptors had more mild phenotypes (Fig. 2C). In addition, the quadruple and quintuple mutants that lacked all three of these receptors (plus other receptor mutations) still had severe deficits in slowing (Fig. 2C). Finally, as described above, quintuple mutants that had any of these three receptors intact displayed NSM-induced slowing, though to different degrees (Fig. 2A-C). This indicates that there are three serotonin receptors whose activation reduces locomotion speed, and concurrent activation of all these receptors produces the slowing response observed in wild-type animals.

The remaining three receptors, *ser-1, ser-5,* and *ser-7,* had modulatory roles in controlling NSM-induced slowing (referred to as the “modulator” receptors). They were not strictly required for serotonin-induced slowing, since the triple mutant lacking all three receptors had no deficit in slowing, compared to wild-type (Fig. 2D). In addition, they could not support robust slowing on their own: the three quintuple mutants with each of these receptors intact were very similar to the sextuple mutant lacking all receptors (Fig. 2A-C). However, in many strains with different combinations of other serotonin receptors deleted, the loss of these receptors caused large deficits in NSM-induced slowing. Fig. 2E shows the effects of deleting each of these receptors in each possible genetic background (displayed as the difference in slowing behavior between a given mutant with the receptor of interest intact versus absent). We found that *ser-7* primarily inhibits slowing, since its deletion most commonly causes animals to display stronger NSM-induced slowing (Fig. 2E, orange bars are mostly negative). *ser-1* (red bars) and *ser-5* (blue bars) had mixed effects (promoting or inhibiting slowing, or having no effect) depending on the background. However, their deletion had little to no effect if the main drivers of slowing (*mod-1, ser-4, and lgc-50*) were already deleted (Fig. 2C), suggesting that they can modulate slowing induced by these receptors. Together with the above analyses, these results suggest that there is one subset of serotonin receptors (*mod-1, ser-4*, and *lgc-50*) whose activation can elicit changes in locomotion behavior. In addition, there is a second subset of receptors (*ser-1*, *ser-5*, and *ser-7*) that, when co-activated together with the other receptors, can modulate the responses.

To discern the exact form of these receptor interactions, we used a modeling approach. Specifically, we constructed a linear model that could predict the amplitude of NSM-induced slowing across all the genotypes, based on which serotonin receptors were present (Fig. 2F). We found that a linear model with six predictor terms, indicating the presence or absence of each receptor, was only able to predict slowing across the genotypes with partial accuracy, even when trained on all the genotypes (Fig. 2G). This indicates that slowing across the full set of genotypes cannot be described as a simple weighted sum of which receptors are present. Therefore, we added additional predictor terms to the model that described the joint presence of two or more receptors (for example, whether *mod-1* and *ser-4* are *both* present), which allows interactions among the receptors to influence the prediction of slowing. We used a procedure where we only added interaction terms if they were statistically required to explain a non-linearity in how two or more receptors interact (in order to favor using the most parsimonious model that fits the data; see Methods). A total of 18 interaction terms were added, which significantly enhanced model performance (5-fold cross-validated performance shown in Fig. 2H; model comparison in Fig. 2I). When this model was trained on data that omitted individual genotypes, it was able to effectively predict the behavior of the genotype withheld from the training data, showing that the way the receptors interacted was similar across genotypes (Fig. S2B). The interaction terms that improved model performance (full list in Fig. S2C) reflected genetic redundancies between specific subsets of receptors, as well as antagonistic interactions, which were primarily between certain “driver” receptors and “modulatory” receptors. These results suggest that extensive interactions between serotonin receptors shape the functional roles of these receptors in modulating behavior.

### Satiety state modulates how serotonin influences behavior by altering the functional roles of the serotonin receptors

An integral aspect of serotonergic signaling in *C. elegans* is its increased functional prominence in fasted animals. Previous studies have shown that when *C. elegans* are fasted they display exaggerated food-induced slowing, an effect referred to as the enhanced slowing response (Sawin et al., 2000). This change is thought to allow animals to increase their exploitation of a food source when they are hungry. The enhanced slowing response is attenuated in animals lacking NSM or serotonin, suggesting that NSM release of serotonin is important for this effect (Iwanir et al., 2016; Rhoades et al., 2019; Sawin et al., 2000). We examined whether NSM-stimulated slowing was exaggerated in animals that had been fasted for three hours compared to well-fed animals. Indeed, the fasted animals displayed more persistent reductions in speed upon NSM stimulation (Fig. 3A-B). This indicates that direct activation of NSM elicits a more robust change in locomotion in fasted animals, recapitulating the enhanced slowing response.

**Figure 3.**
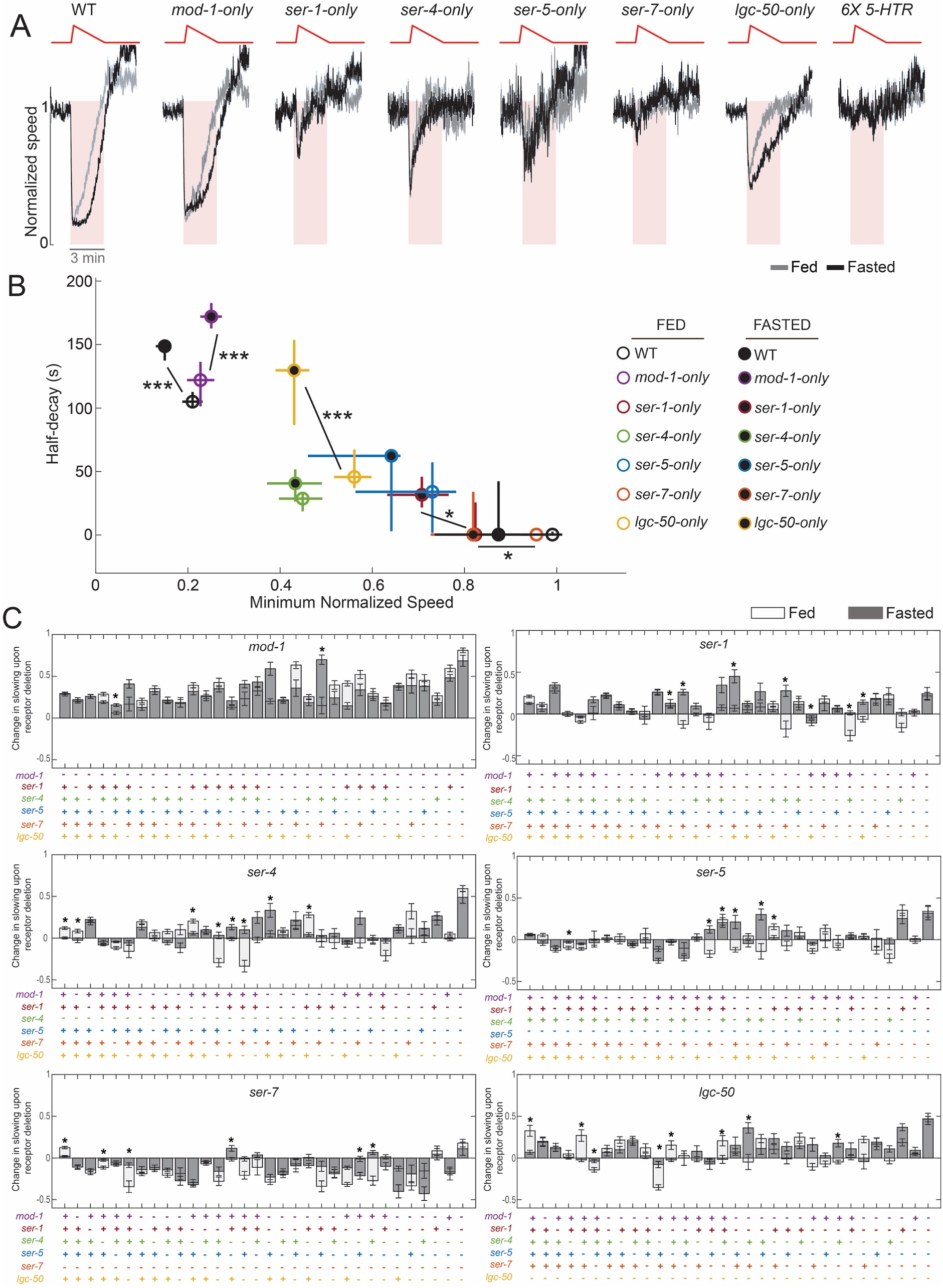
Satiety impacts how the serotonin receptors control locomotion. (A) Mean speed traces of animals of the indicated genotypes in response to NSM::Chrimson activation with the indicated waveform of light (top, red; medium intensity stimulus described in Fig. 2). As indicated, animals were either well-fed or fasted for three hours. Data are shown as means ± SEM. Data are quantified in panel (B). (B) Quantification of data in panel (B), showing level of slowing (x-axis) and decay of speed back to baseline (y-axis) for each genotype. N=44-495 animals per condition. *p<0.05, ***p<0.001, level of slowing in fed versus fasted, Bonferroni-corrected empirical p-value that difference between distributions of either level or decay of slowing is non-zero. (C) Six plots depicting the change in NSM-induced slowing that is caused by a given receptor’s mutation, in the different genetic backgrounds (i.e. combinations of other serotonin receptor mutations) that were tested. Data are shown as in Fig. 2I, except that here the data from fasted animals is also shown, as indicated. Y-axis values reflect the difference in the level of slowing in the genotype indicated, compared to a corresponding genotype that is the same except the receptor in question is present (see illustrative cartoon depicting this at top of Fig. 2E). *p<0.05, Bonferroni-corrected empirical p-value that the y-axis values (effect of receptor deletion) are different between fed and fasted. In addition, see Main Text for additional criteria to consider a genotype as being significant here. Average n across all the genotypes is 278 animals for fed and 328 animals for fasted. Data are shown as means ± SEM.

In principle, fasting could enhance NSM-stimulated slowing by (i) increasing NSM-stimulated serotonin release, (ii) increasing serotonin receptor expression; (iii) enhancing how the receptors drive slowing due to changes in receptor-expressing cells (for example, stronger coupling of those cells to behavioral outputs), or (iv) altering how the receptors interact to drive slowing (for example, making receptors act more synergistically). To begin to distinguish among these possibilities, we examined whether slowing was enhanced by fasting in each of the quintuple mutants that have one serotonin receptor remaining (Fig. 3A-B). We found that NSM-stimulated slowing was enhanced by fasting in animals with *mod-1* or *lgc-50* intact, but not in animals with only *ser-4* left intact (the other quintuple mutants had minimal slowing in the fed state, so they are hard to interpret). This suggests that fasting enhances the locomotion change induced by activation of some, but not all, serotonin receptors. This observation renders it unlikely that this effect can be explained by an increase in serotonin release upon optogenetic NSM stimulation in fasted animals, as this would have been predicted to increase NSM-stimulated responses in the *ser-4*-only mutant also. (Critical to this interpretation is the observation that the level of slowing in well-fed *ser-4*-only animals is increased in a dose-dependent manner by increasing the intensity of the optogenetic stimulus, and yet, at intermediate levels of stimulation, fasting still did not enhance slowing; Fig. S3A-B). The enhanced slowing mediated by certain receptors does not appear to be due to changes in the expression of those receptors. We found that the neurons in which the receptors are expressed did not change with fasting (see Fig. 5 below). In addition, the overall levels of mRNA translation for each of the six receptors in well-fed versus fasted animals were not significantly different (based on our previous ribotagging results in McLachlan et al., 2022, which used animals of the same age and the same fasting duration). This suggests that the impact of MOD-1 and LGC-50 receptor activation on neural circuit activity may be enhanced by fasting.

To examine whether fasting also alters the functional interactions between the receptors, we recorded NSM-stimulated behavioral responses in well-fed and fasted conditions for all 64 serotonin receptor mutant strains. We asked whether the effect of deleting a given serotonin receptor in each possible genetic background (i.e. mutant background with a specific subset of serotonin receptors already deleted) was different in well-fed versus fasted animals (Fig. 3C). We only considered there to be a difference between fed and fasted conditions if three criteria were met: (1) the loss of the receptor led to a different effect on behavior in well-fed and fasted animals (enhanced vs attenuated vs no-effect on NSM-stimulated locomotion); (2) the magnitude of the behavioral change due to loss of the receptor was significantly different between the fed and fasted conditions (since criterion (1) could be susceptible to p-values changing from 0.04 to 0.06, etc) ; and (3) these differences could be observed under conditions where neither parental group (i.e. the strain with the given serotonin receptor still intact) had saturating levels of slowing, since this would complicate interpretation of lack of an effect upon deletion of a receptor. Using these criteria, we observed many significant differences between fed and fasted animals. These differences commonly involved a given receptor deletion having no effect in one satiety state and a strong effect in the other state. For example, whereas the role of *ser-7* was similar in both states, *ser-1* and *lgc-50* had stronger contributions to slowing in the fasted state. In addition, there were clear examples of inversions of function, where loss of the receptor exaggerated slowing in well-fed animals, but attenuated slowing in fasted animals (in particular, for *ser-5*). These results suggest that interactions among the receptors are modulated by the animal’s satiety state. Taken together, these results suggest that the enhanced responses to serotonin release in fasted animals are due to enhanced effects of *mod-1* and *lgc-50* on downstream circuits and functional changes in interactions among the receptors.

### The MOD-1 and SER-4 receptors confer behavioral responses to different patterns of serotonin release, due to their distinct sites of expression

In analyzing the serotonin receptor mutants’ behavioral phenotypes, we observed that different serotonin receptors exhibited striking differences in terms of the behavioral dynamics that they support. This is readily apparent in the quintuple mutants that have only one serotonin receptor intact (Fig. 4A-B shows a direct comparison). For the three “driver” receptors that can drive slow locomotion on their own, the maintenance of the slow locomotion during continued NSM stimulation differed dramatically. Whereas *mod-1*-only mutants displayed reduced locomotion as long as NSM was active, the *ser-4*-only mutants only displayed a transient reduction in locomotion speed at stimulation onset that rapidly decayed. *lgc-50*-only animals displayed an intermediate decay rate (Fig. 4A-B). In addition, in contrast to *mod-1*-only animals, *ser-4*-only animals did not show different responses to short versus long optogenetic stimuli (Fig. 4C-D). This suggests that *mod-1*-only animals can track the continued release of serotonin, whereas *ser-4*-only animals cannot. These effects could not be explained as an indirect consequence of different magnitudes of slowing, since these differences in decay rates were consistent across several different NSM stimulation intensities that evoked different levels of slowing. We also asked whether these differences observed in the quintuple mutants generalized to the other genotypes that we recorded. Indeed, all genotypes with *mod-1* and *lgc-50* intact had sustained NSM-induced slowing, and all mutants lacking these two receptors did not, suggesting that these two receptors are critical for sustained slowing (Fig. 4E). These data suggest that activation of different serotonin receptors drives different dynamical changes in behavior.

**Figure 4.**
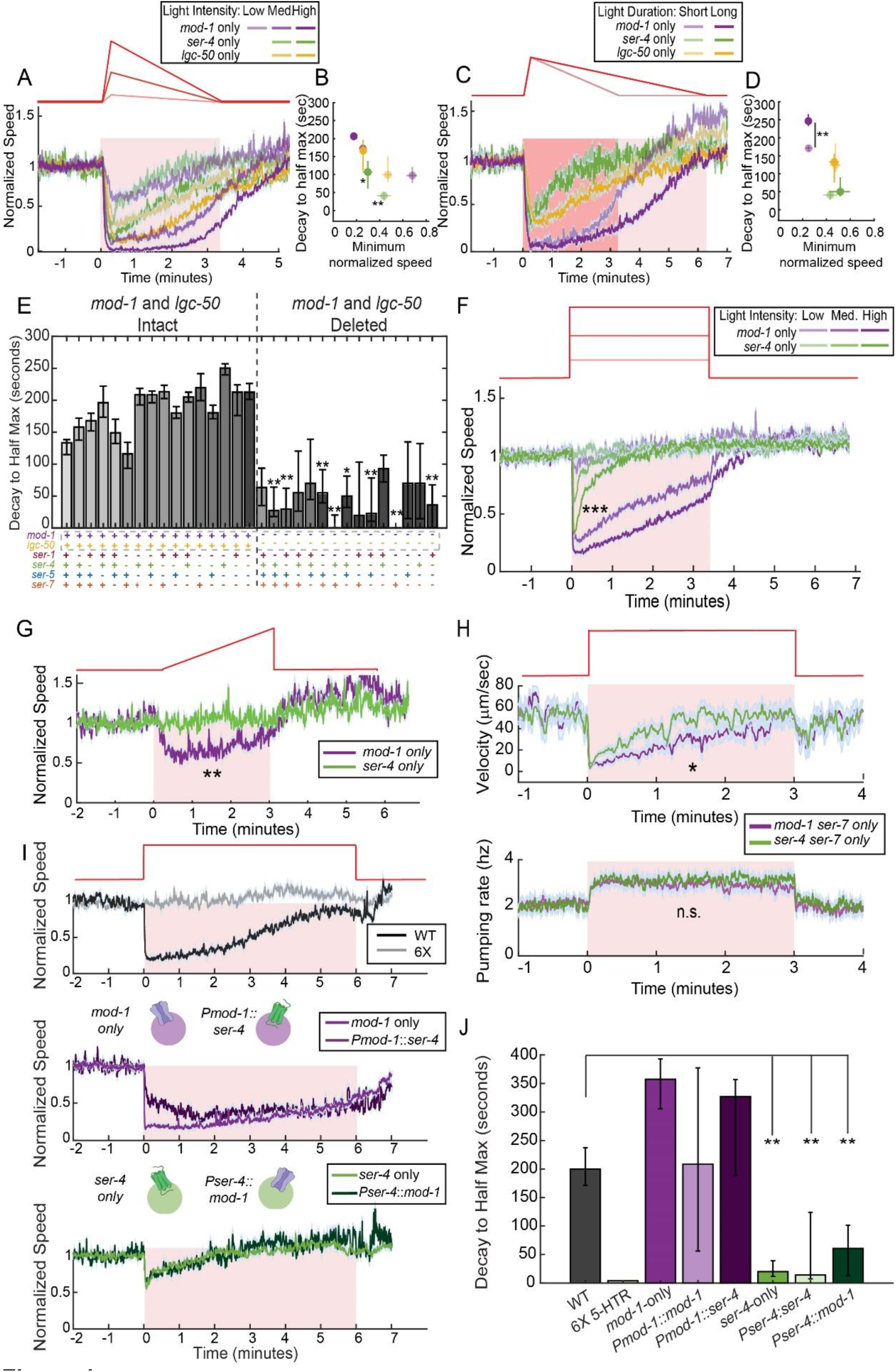
The *ser-4* and *mod-1* serotonin receptors mediate responses to different patterns of serotonin release, due to their distinct sites of expression in the connectome. (A) Changes in animal speed induced by NSM::Chrimson activation with the indicated waveforms of light (red, top). For *mod-1*-only, we show the lowest intensity stimulus, but this is not shown for the other two mutants since slowing is negligible for them at that low stimulus intensity. Data are shown as means ± SEM. Data are quantified in panel (B). (B) Quantification of traces shown in panel (A). **p<0.01, *ser-4-*only medium intensity stimulation versus all other conditions, Bonferroni-corrected empirical p-value that difference between distributions of decays is non-zero. *p<0.05, *ser-4*-only high intensity stimulation versus *lgc-50*-only high intensity and *mod-1*-only medium and high intensity, Bonferroni-corrected empirical p-value that difference between distributions of decays is non-zero. N=116-495 animals per condition. Dot are means and error bars show ± 95% confidence interval. (C) Changes in animal speed induced by NSM::Chrimson activation with the indicated waveforms of light (red, top). Data are shown as means ± SEM. Data are quantified in panel (D). (D) Quantification of traces shown in panel (C). **p<0.01, medium-versus high-intensity stimulation conditions for a given genotype, Bonferroni-corrected empirical p-value that difference between distributions of decays is non-zero. N=103-495 animals per genotype. Dots are means ± 95% confidence interval. (E) Speed decay rates back to baseline after maximal slowing, for the indicated genotypes. Genotypes are indicated as dots along the x-axis, where the presence of a dot indicates that the gene is intact (rather than mutated). *p<0.05, **p<0.01, WT versus indicated genotype, Bonferroni-corrected empirical p-value that difference between distributions of decays is non-zero. Average n=284. Data are shown as means ± 95% confidence interval. (F) Changes in animal speed induced by NSM::Chrimson activation with the indicated waveforms of light (red, top). Data are shown as means ± SEM. ***p<0.001, max intensity ser-4-only condition versus medium-intensity *mod-1*-only condition since they are most closely matched for level of slowing, empirical p-value that difference between distributions of decays is non-zero. N=356-438 animals per condition. (G) Changes in animal speed induced by slow-ramping NSM::Chrimson activation, as indicated. Data are shown as means ± SEM. **p<0.01, empirical p-value that difference between distributions of slowing magnitude is non-zero. N=177-466 animals per condition. (H) Changes in velocity and pumping rates for animals of the indicated genotypes, elicited by a single three minute block stimulation of NSM::Chrimson. *p<0.05, Bonferroni-corrected t-test of lights-on behavior between the two genotypes. n.s., not significant. N=84-96 stimulation events per condition (three trials per animal). Data are shown as means ± SEM. (I) Changes in animal speed induced by NSM::Chrimson activation in the indicated genotypes. Inset cartoons depict the receptor swap transgenic lines. Data are shown as means ± SEM. Data are quantified in panel (J). (J) Quantification of data in (I). Decay rates back to baseline speed, after maximal slowing, for animals of the indicated genotypes. **p<0.01, WT vs. indicated genotypes, Bonferroni-corrected empirical p-value that difference between distributions of decays is non-zero. N=47-306 animals per genotype. Data are shown as means ± 95% confidence interval.

**Figure 5.**
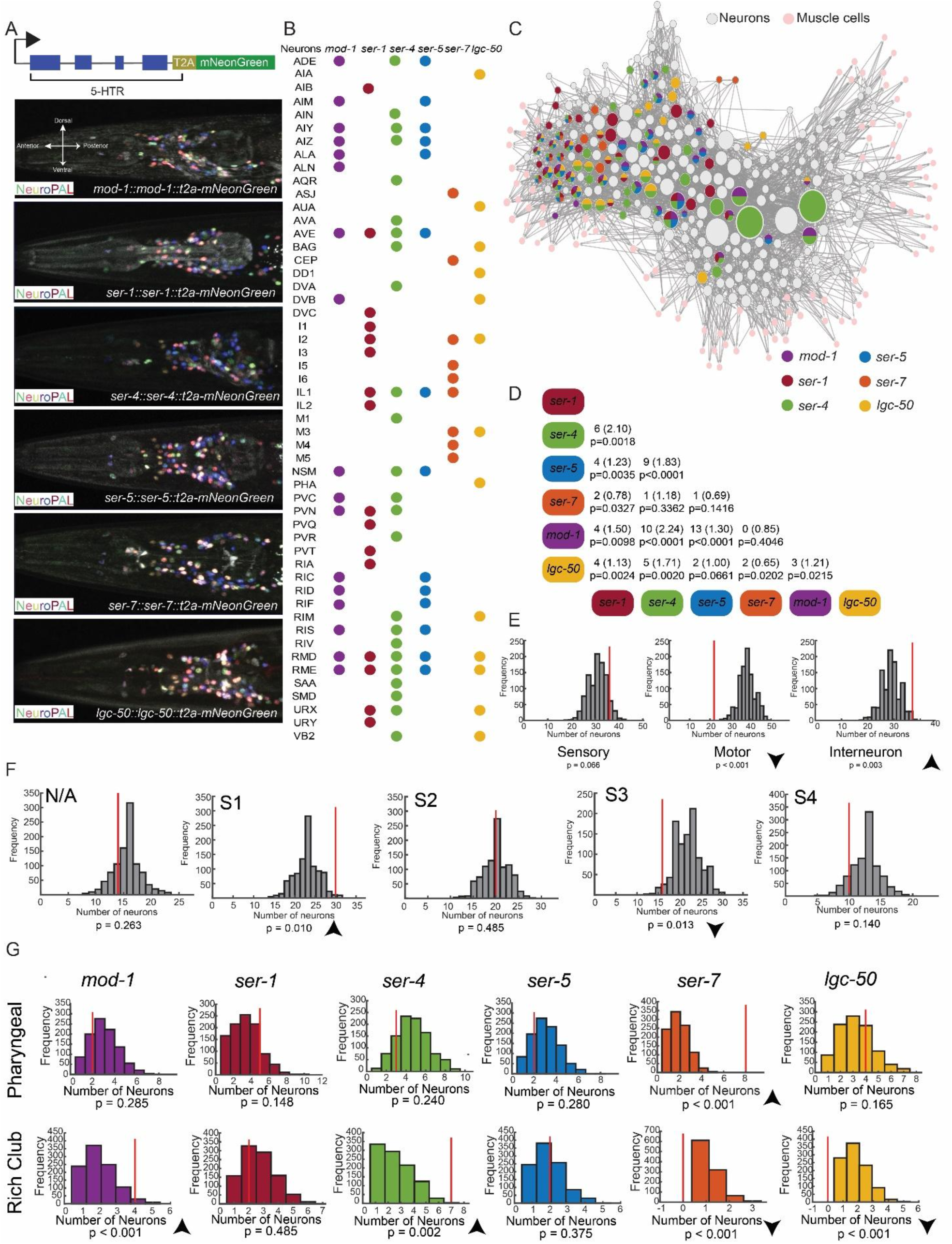
Mapping all sites of serotonin receptor expression across the *C. elegans* connectome. (A) Top: Cartoon illustrating how CRISPR/Cas9 editing of a serotonin receptor gene was used to introduce a soluble mNeonGreen reporter for a given serotonin receptor’s expression. Six such lines were constructed, one for each serotonin receptor gene. Below: Example images depicting mNeonGreen reporter expression for each serotonin receptor gene (white) and NeuroPAL labels (colors) that allow us to infer neural identity. (B) A list of which serotonin receptor gene reporters were detected in which classes of neurons. Dots indicate that expression was detected. Neurons without any detected serotonin receptor reporter expression are not listed here. (C) Diagram of the *C. elegans* wiring diagram, with colors illustrating which receptors are expressed in which neurons. Adjacency of neurons in this plot indicates interconnected wiring so that the plot roughly depicts the wiring organization of the connectome. (D) Chart showing the number of neurons that show overlapping expression for pairwise combinations of serotonin receptors. For each receptor pair, we show the number of neurons with overlapping expression and, in parentheses, the number that would have been expected by chance. In addition, the multiple comparison-corrected p-value is for the permutation test for overlap, described in Methods. (E) Enrichment of serotonin receptor expression (pooling all receptors) in sensory neurons, interneurons, and motor neurons. The red line indicates the actual number of receptors observed in the group of cells and the gray histogram indicates the numbers that would be expected at random (randomly sampled 1000 times). The multiple comparison-corrected p-value is then the result of a permutation test, described in Methods. (F) Enrichment of neurons in the different strata of the *C. elegans* nerve ring, shown as in (E). Segmentation of the *C. elegans* neurons into defined strata is based on previous work (Moyle et al., 2021). N/A refers to neurons not categorized into any of the four strata. Multiple comparison-corrected p-values are shown. (G) Enrichment of each of the serotonin receptors in the Rich Club neurons (Towlson et al., 2013) and the pharyngeal nervous system, shown as in (E) except splitting out each serotonin receptor. Multiple comparison-corrected p-values are shown.

The difference in the behavioral dynamics supported by *mod-1* and *ser-4* was especially striking, suggesting that *mod-1* conferred behavioral responses to continued NSM activation, while *ser-4* only conferred responses to the onset (or high positive derivative) of NSM activation. To more closely examine this, we compared these strains’ behavioral responses to other temporal patterns of NSM activation. A stimulation pattern with an immediate activation of NSM::Chrimson further accentuated this difference: *ser-4*-only animals transiently slowed down upon NSM activation, while *mod-1*-only animals maintained slow locomotion (Fig. 4F). We also examined responses to an NSM stimulation pattern with a slow, ramping onset that never contained a high positive derivative. *mod-1*-only animals slowed in response to this optogenetic stimulus. However, *ser-4*-only animals did not respond at all, further confirming that this receptor only confers a response to a sharp increase in NSM activation (Fig. 4G).

We wanted to examine whether these findings generalize to natural stimuli that evoke NSM activation and, more broadly, examine how *mod-1* and *ser-4* contribute to transient and persistent food-driven behavioral changes. Thus, we examined mutant animals’ speed upon bacterial food patch encounter, which leads to immediate and robust activation of NSM (Fig. S4; (Iwanir et al., 2016; Rhoades et al., 2019)). Food-induced slowing is partially dependent on serotonin, so animals lacking all six serotonin receptors displayed an attenuated speed reduction at all time points after food encounter, though as described above loss of serotonin signaling only partly attenuates slowing (Fig. S4A, gray trace). *mod-1*-only animals had wild-type slowing, matching the optogenetics results. In addition, *ser-4*-only animals displayed a rapid slowdown upon food encounter that exceeded the level observed in the sextuple mutants, but after one minute their speed matched the sextuple mutants, suggesting that the presence of *ser-4* induces slowing immediately after food encounter, but has no effect at later timepoints. This also provides a qualitative match to the optogenetic results. Together, these data suggest that *mod-1* confers sustained behavioral responses to sustained NSM activation, while *ser-4* confers responses to the onset of NSM activation.

*mod-1* is a serotonin-gated chloride channel (a 5HT3 homolog) and *ser-4* is a Gi/o-coupled G protein-coupled receptor (a 5HT1A homolog). Previous work has suggested that both act in an inhibitory fashion (Olde and McCombie, 1997; Ranganathan et al., 2000). In principle, the difference in the locomotion dynamics of the quintuple mutants with only *mod-1* or *ser-4* could be caused by differences in serotonin release dynamics (due to feedback to NSM, etc), the molecular properties of the receptors, or the properties of the neurons where these inhibitory receptors are expressed (for example, their extent of recurrent connectivity that could shape neurons’ ongoing dynamics). To distinguish among these possibilities, we first examined whether altered serotonin release dynamics could explain these effects. If serotonin release dynamics are different in these two mutants, then other NSM-induced behavioral changes should also show different dynamics. We tested this by examining the dynamics of NSM-induced feeding behavior in mutant animals with *mod-1* or *ser-4* present (along with *ser-7*, the serotonin receptor required for serotonin-induced feeding) (Hobson et al., 2006). The difference in locomotion decay rates was still clearly evident when comparing *mod-1/ser-7*-only animals to *ser-4/ser-7*-only animals (Fig. 4H). However, the dynamics of NSM-stimulated feeding behavior were indistinguishable when comparing the two strains (Fig. 4H; feeding behavior was not saturated by NSM stimulation, as we found that these mutants with *ser-7* intact pump at ∼5Hz in the presence of food, similar to wild-type). This result indicates that the difference in behavioral dynamics supported by *mod-1* and *ser-4* does not extend to other serotonin-induced behavioral changes, suggesting it is unlikely that serotonin release has been affected.

We next tested whether these functional differences between *mod-1* and *ser-4* were due to the different cell types in which they were expressed or, alternatively, their different molecular properties. Given that they are both inhibitory receptors, we tested this by swapping their sites of expression and examining the impact on NSM-induced behavioral responses. These experiments were conducted via transgenic rescue in the sextuple mutant background. Sextuple mutant animals expressing *mod-1* under the *ser-4* promoter (*Pser-4::mod-1*) displayed transient slowing upon serotonin release, matching the *ser-4*-only mutant and the *Pser-4::ser-4* transgenic rescue animals (Fig. 4I-J). In contrast, animals expressing *ser-4* under the *mod-1* promoter (*Pmod-1::ser-4*) displayed maintained slowing throughout the optogenetic stimulus, matching the *mod-1*-only mutants and *Pmod-1::mod-1* animals (Fig. 4I-J). Thus, the functional roles of the receptors tracked the promoter used, rather than the molecular identity of the receptor being expressed. We note that the expression patterns of these promoters might only partially recapitulate the native expression patterns of these genes. Nevertheless, these results provide direct evidence that the sites of expression of these two receptors, rather than their molecular properties, play a dominant role in conferring the kinetics of the NSM-induced behavioral response that they support. The *ser-4*-expressing neurons are sensitive to the positive derivative of NSM activity, while the *mod-1*-expressing neurons are sensitive to the overall level of NSM activity.

### Mapping sites of serotonin receptor expression across the *C. elegans* connectome

The above results indicate that the cellular expression patterns of the serotonin receptors are important in conferring their functional roles in controlling behavior. Therefore, we next sought to determine where each serotonin receptor was expressed across the connectome. To do so, we used CRISPR/Cas9 genome editing to insert T2A-mNeonGreen fluorescent reporter cassettes into each of the six serotonin receptor genes, just before their stop codons (Fig. 5A). Under this strategy, the T2A self-cleaving peptide allows mNeonGreen to be expressed as a cytosolic protein. The expression of mNeonGreen in each of these six strains should match the native pattern of each serotonin receptor’s expression, since it should be under similar transcriptional and translational control. We chose to perform this mapping, as opposed to relying on existing single-cell sequencing data (Taylor et al., 2021), because we wanted to assess functional levels of expression in well-fed and fasted one-day old adult animals.

To determine the ground-truth identities of the fluorescently labeled neurons in each of these six strains, we crossed the NeuroPAL transgene into each of the mNeonGreen reporter strains (Fig. 5A; Fig. S5A). The NeuroPAL transgene expresses three fluorescent proteins (NLS-BFP, NLS-OFP, NLS-mNeptune) under well-defined genetic drivers, so that neural identity can be easily inferred from position and multi-spectral fluorescence (Yemini et al., 2021). For each reporter strain, we annotated the identities of all mNeonGreen-positive neurons in 5-10 adult animals. Results were highly consistent across animals and were nearly identical for well-fed and fasted animals (Fig. S5B). Consistent with this, our previous ribotagging data in well-fed versus fasted animals showed no change in expression of the six receptors (McLachlan et al., 2022; though we cannot rule out more subtle changes in the levels of expression in subsets of cells). Each receptor was expressed in at least 8 of the 118 *C. elegans* neuron classes, with a range of 8-24 neuron classes per receptor (Fig. 5B). Altogether, 52 neuron classes (44% of all neuron classes) expressed at least one serotonin receptor (Fig. 5C). Our results were broadly in agreement with existing single-cell sequencing of FACS-sorted cells from L4 animals (Taylor et al., 2021), though there were differences (both cells detected via sequencing, but absent here; and cells detected here, but absent in the sequencing data).

We performed several analyses on these expression data to examine how the serotonin receptors were distributed across the connectome. First, we examined the similarity of the expression patterns of the receptors. While each serotonin receptor was expressed in a unique pattern, the level of overlap between the receptors was higher than expected by chance for most of the pairs (Fig. 5D). This resulted in some individual neuron classes expressing as many as five different serotonin receptor genes. Importantly, we used an identical strategy to construct T2A-mNeonGreen reporters for several other GPCRs (in the olfactory receptor gene family) that showed completely different expression patterns, indicating that correlated expression among the serotonin receptors is not an artifact of the reporter strategy (McLachlan et al., 2022). This high incidence of co-expression of the serotonin receptors could plausibly underlie some of the functional interactions among the receptors.

We also considered which cell types and circuits in the connectome expressed the serotonin receptors. The receptors displayed significantly enriched expression in interneurons, and significantly de-enriched expression in motor neurons (Fig. 5E). Their expression in sensory neurons matched the level expected by chance (Fig. 5E). This suggests that serotonin primarily modulates interneurons and acts upstream of the motor neurons of *C. elegans*. The *C. elegans* nerve ring has been described to consist of four major strata or interconnected groups of neurons with different functional roles (Brittin et al., 2021; Moyle et al., 2021). Serotonin receptor expression was enriched in stratum 1, which is associated with head movements and exploration, and de-enriched in stratum 3, associated with aspects of movement control (Fig. 5F). The *mod-1* and *ser-4* receptors also displayed enriched expression in the “Rich Club” neurons, a smaller set of densely connected hub interneurons that span the strata and exert a strong influence on locomotion (Fig. 5G) (Towlson et al., 2013). Finally, *ser-7*, but not the other receptors, was enriched in the pharyngeal nervous system (Fig. 5G), consistent with its prominent role in controlling serotonin-induced feeding (Hobson et al., 2006). Together, these analyses reveal broad and correlated expression of the serotonin receptors in circuits that control exploration and locomotion.

### Different temporal phases of the MOD-1-induced behavioral change can be mapped onto different sites of expression

Having established the expression patterns of the serotonin receptor genes, we next performed cell-specific rescue experiments to determine which sites of receptor expression are functionally important for locomotion. We chose to examine this in detail for the *mod-1* receptor, since sole expression of *mod-1* can confer rapid and sustained changes in locomotion. To perform these rescues, we engineered a mutant allele of the native *mod-1* gene in which the orientation of several exons was inverted, inactivating the gene (Fig. 6A). These exons were flanked by dual lox sites that could invert the orientation of the exons upon Cre expression, which would be predicted to restore gene function in Cre-expressing cells. Importantly, this strategy constrains expression levels so that cell-specific expression of *mod-1* never exceeds the native level of expression. This allele was engineered into the *mod-1*-only quintuple mutant, so that we could perform cell-specific rescues in a mutant background with no other functional serotonin receptors. Indeed, we found that this strain displayed no serotonin-induced slowing, matching the sextuple mutant lacking all receptors (Fig. 6B). This indicates that inversion of the *mod-1* exons successfully inactivated *mod-1* function. We then expressed Cre in this strain using a pan-neural promoter (*tag-168*) and found that this led to a full rescue of the locomotion phenotype (Fig. 6B), indicating that Cre expression can successfully invert the *mod-1* gene in neurons and rescue *mod-1* function.

**Figure 6.**
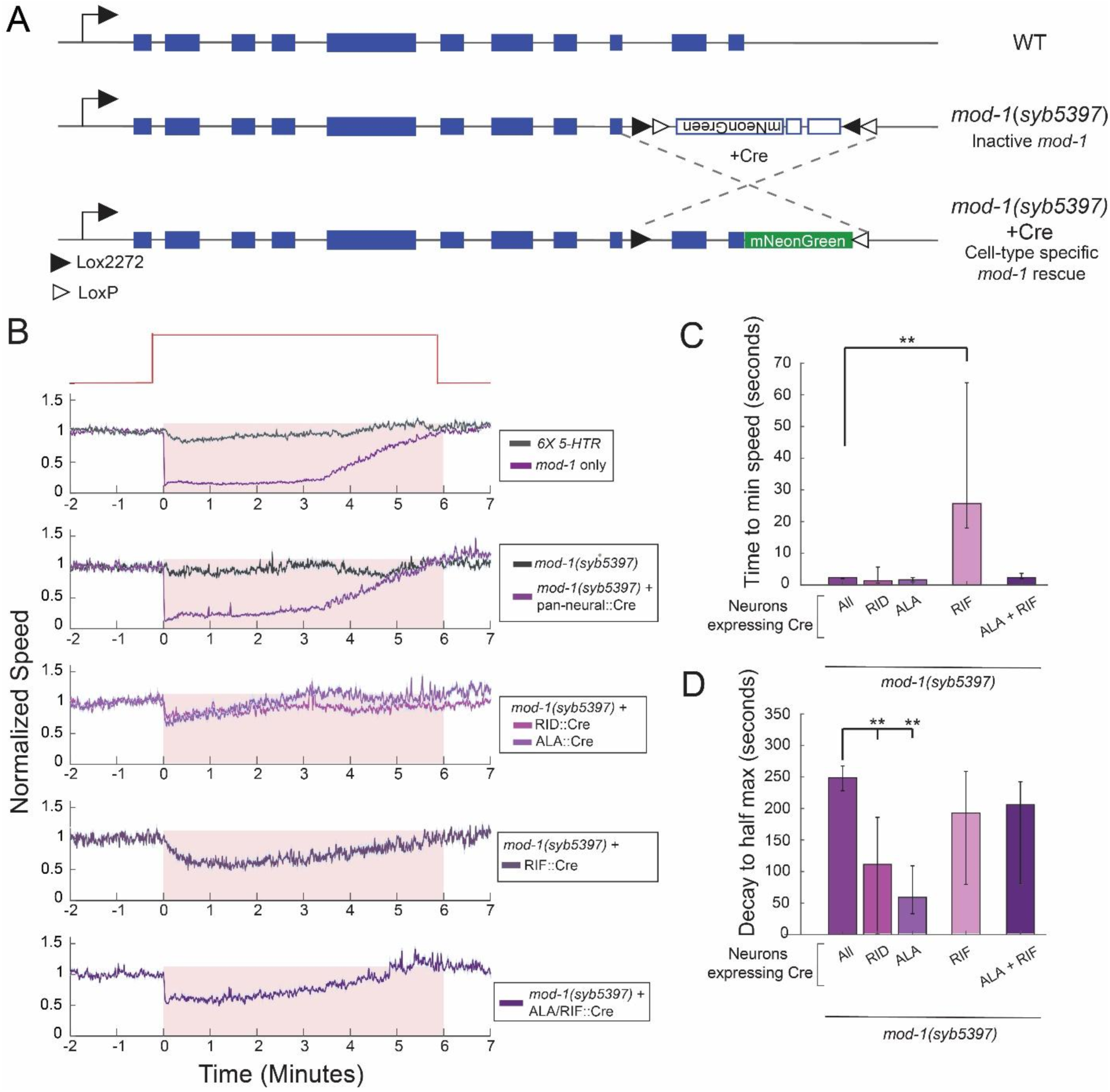
The different temporal phases of the *mod-1*-induced behavioral change are supported by *mod-1* function in different neuron types. (A) Cartoon illustrating the genetic strategy for *mod-1* rescue experiments. CRISPR/Cas9 gene editing was used to edit the endogenous *mod-1* gene (top). This results in a *mod-1* allele that is inactive (middle) until Cre expression inverts the exons back to their active orientation (bottom). (B) Changes in animal speed induced by NSM::Chrimson activation with the indicated waveform of light (top, red), shown for the indicated genotypes. Data are shown as means ± SEM, and are further quantified in panels (C) and (D). (C) Quantification of data in (B). Latency to time point of maximal slowing upon NSM::Chrimson activation, shown for the indicated genotypes. Promoters used for Cre expression were: *Ptag-168* (pan-neuronal), *Pflp-14* (RID), *Pver-3* (ALA), *Podr-2b* (RIF). **p<0.01, pan-neuronal Cre versus indicated cell-specific rescue line, Bonferroni-corrected empirical p-value that difference between distributions of latencies is non-zero. N=41-157 animals per condition. Data are shown as means ± 95% confidence interval. (D) Quantification of data in (B). Decay rate back to baseline after maximal slowing, shown for the indicated genotypes. Promoters used for Cre expression were: *Ptag-168* (pan-neuronal), *Pflp-14* (RID), *Pver-3* (ALA), *Podr-2b* (RIF). **p<0.01, pan-neuronal Cre versus indicated cell-specific rescue line, Bonferroni-corrected empirical p-value that difference between distributions of decays is non-zero. N=41-157 animals per condition. Data are shown as means ± 95% confidence interval.

We next performed cell-specific rescues in the majority of the *mod-1*-expressing neurons, using a panel of Cre drivers that drive expression in unique sets of these neurons. We identified three functionally distinct classes of cells. First, there were several neurons where *mod-1* genetic rescue did not restore slowing at all (RME, RIS, ADE, NSM, RIC, AIM, AIY, AIZ, AVE; Fig. S6A). Second, there were two neurons where *mod-1* genetic rescue caused animals to display a transient slowdown upon NSM activation, similar to what was observed in *ser-4*-only animals (RID, ALA; Fig. 6B-D). Third, there was one neuron where *mod-1* rescue led to a slow onset, sustained slowing upon NSM activation (RIF, Fig. 6B-D). This latter phenotype with an exceptionally slow onset had not been observed in any of the genetic mutants studied above. Interestingly, we found that we could restore both the transient and sustained slowdown by combined genetic rescue in cells from both categories (ALA+RIF; Fig. 6B-D). These results suggest that the initial slowdown upon NSM activation and the sustained slowdown that continues during NSM activation are separable effects, mediated by *mod-1* function in different neurons. Distinct temporal phases of the behavioral response to serotonin release can be mapped onto different neurons. In addition, given the partial rescues observed for all of these Cre lines, these results further suggest that *mod-1* likely functions in many neurons to promote slowing. In order to fully rescue *mod-1* function, expression may need to be restored in many neurons, each with unique contributions.

### NSM activity is associated with widespread changes in neural activity across the brain as animals freely move

The above results suggest that serotonin release from NSM activates several different serotonin receptor types in distinct neuron classes to modulate locomotion. We next sought to examine how NSM’s ongoing activity was associated with changes in neural activity in these distributed circuits. Therefore, we performed brain-wide calcium imaging in freely-moving animals as they navigated to and encountered a food patch, a natural context in which NSM is activated to modulate locomotion. For these recordings, we used a strain expressing NLS-GCaMP7f (a calcium sensor) and NLS-mNeptune2.5 (a red fluorescent protein) in all *C. elegans* neurons (Fig. S7A; Atanas et al., 2022). In addition, NLS-TagRFP-T is expressed in NSM and a few other neurons, allowing us to identify NSM in these recordings. Freely-moving animals were recorded on a previously-described, live-tracking microscope with two light paths (Atanas et al., 2022). A spinning disc confocal below the animal allows for fast volumetric imaging of fluorescent neurons in the head. Above the animal, a low-magnification light path captures images for behavior quantification (example images in Fig. S7A). A live-tracking system keeps the animal centered in view as it freely moves. We used our recently-described automated data processing pipelines to extract GCaMP traces and behavioral variables from these videos, which we previously confirmed faithfully extracts calcium traces from single neurons over time (Atanas et al., 2022). In addition, we previously recorded animals expressing pan-neuronal NLS-GFP and NLS-mNeptune to estimate motion artifacts in data recorded on this imaging platform and found that they are negligible (Atanas et al., 2022). Nevertheless, we use GFP recordings to correct and control for any small motion artifacts in all analyses below (see Methods). These tools allow us to record brain-wide activity and behavior as animals freely navigate to and encounter a food patch.

We recorded data from seven animals over 16 minutes as they navigated towards and eventually encountered a bacterial food patch (Fig. 7A). Animals were fasted for three hours prior to the recording in order to make the role of serotonin in behavioral control more prominent (Rhoades et al., 2019; Sawin et al., 2000). As expected, animals abruptly slowed down upon food encounter (Fig. 7B-C). In addition, their oscillatory head movements slowed and their feeding rates increased (Fig. 7B). NSM was inactive while animals were off food, but its activity sharply increased upon food encounter and it showed phasic bouts of activity while animals moved and ate (Fig. 7B, bottom). We examined whether NSM’s ongoing activity was correlated with ongoing changes in speed, head bending, and feeding. To do so, we examined the correlation of the NSM GCaMP signal to these behavioral variables when animals were on food. We favored this approach over an analysis of all pre- and post-food-encounter time points, because NSM is activated upon food encounter and an analysis across all time points would inevitably lead to the conclusion that NSM is associated with all food-induced behaviors, even if such relationships are not convincingly time-locked. Indeed, we found that NSM activity showed a significant negative correlation with speed and the rate of head movement, indicating that time-varying increases in NSM activity were associated with reduced speed and head movement (Fig. 7H; see Methods for a description of shuffle controls and statistics). NSM activity was also frequently associated with increased feeding. This suggests that under our recording conditions, animals display NSM-associated reductions in speed and head movements and an increase in feeding when they encounter food.

**Figure 7.**
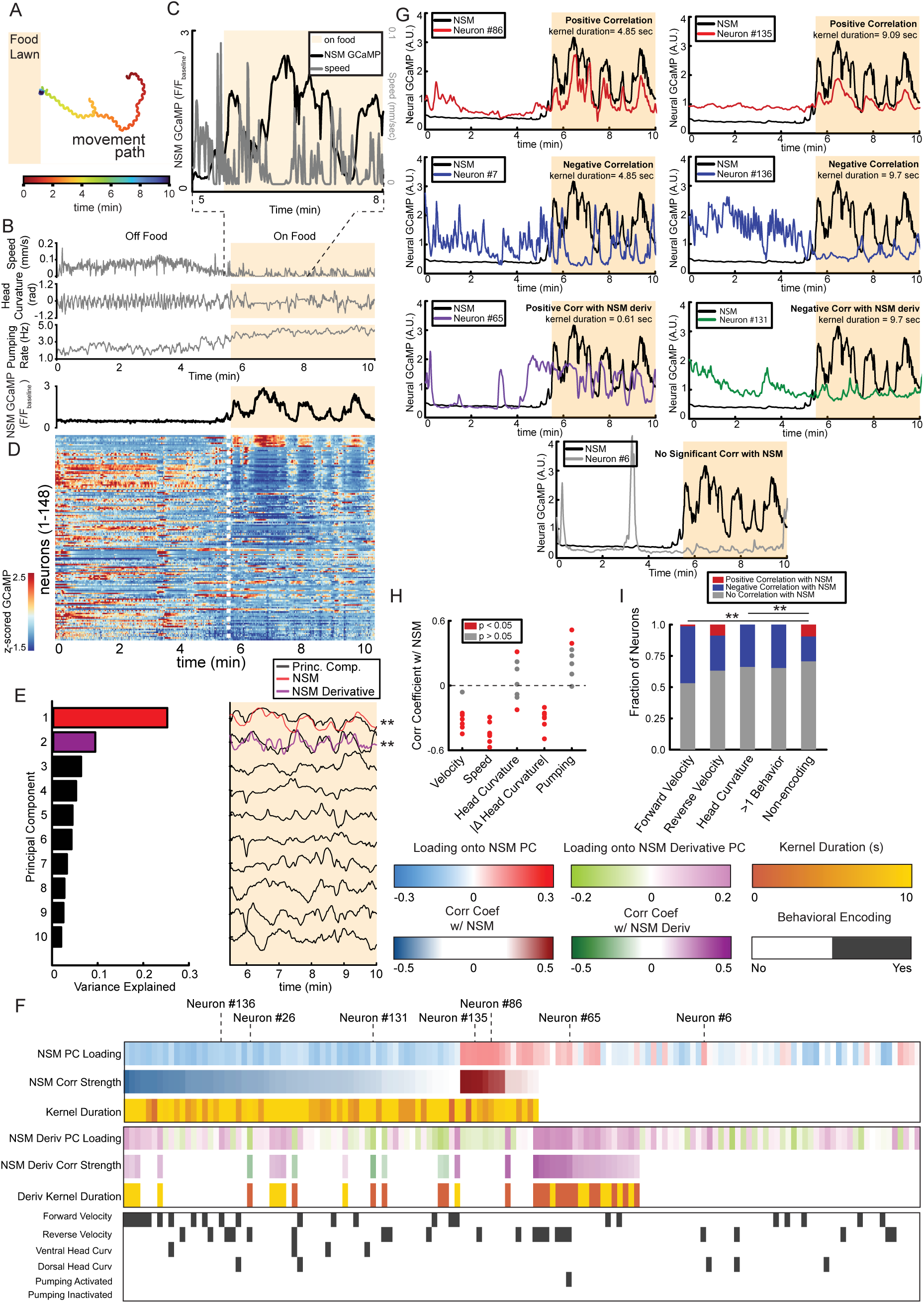
NSM activity is associated with widespread changes in brain-wide dynamics in freely-moving animals. (A) Example animal movement path during a freely-moving brain-wide calcium imaging recording. Animals were placed on an agar pad in a region where there was no food and they moved freely until they found the *E. coli* OP50 food patch. Brain-wide calcium imaging and behavioral recordings were ongoing during this entire time (pre- and post-food-encounter). This single animal’s data is analyzed in panels (A-G) of this figure as an illustrative example, and data across animals are summarized in in panels (H-I) and by statistics cited in the Main Text. (B) Example behavioral and NSM GCaMP data from an animal recorded during freely-moving food patch encounters. The beige background color indicates when the animal encounters the food patch. (C) Zoomed in detail of a portion of the data in panel (B) showing synchronous NSM GCaMP and animal speed during a time period after the food encounter. Note the clear inverse relationship between NSM GCaMP and animal speed. (D) Heatmap depicting calcium traces from 148 simultaneously recorded neurons from the freely-moving animal shown in (A-G). Note that a large fraction of neurons (primarily at the top) show dynamics associated with NSM (black trace directly above). (E) An analysis of the principal components from this example, which reveals the main modes of neural dynamics in this example animal. PCA was conducted using the post-food-encounter time epoch of the dataset. Left: variance explained by each of the principal components. Right: each principal component shown over time. Note that PC1 is correlated to NSM and PC2 is correlated to the derivative of NSM (**p<0.01, shuffle test for significant correlation, see Methods). (F) An analysis of all the neurons recorded in the example dataset depicted in panels (A-G). Each neuron is shown as a column and the following descriptors of information are provided for each neuron, in the rows: (1) NSM PC loading: the factor loadings for PC1 from this dataset. (2) NSM Corr Strength: the correlation coefficient of the neuron with its optimal kernel-convolved NSM (see text and Methods). No value is shown if the correlation was not significant. (3) The duration of the optimal kernel from (2), which provides information about the timescale over which the neuron correlates with NSM. (4) NSM Deriv PC Loading: the factor loading for PC2 from this dataset. (5) NSM Deriv Corr Strength: the correlation coefficient of the neuron with its optimal differentiator-kernel-convolved NSM (see text and Methods). No value is shown if the correlation was not significant. (6) The duration of the derivative kernel, which provides information about the timescale of the correlation between the neuron and NSM’s derivative. (7) Indicators of whether the neuron encoded any of the listed motor programs during the pre-food-encounter time epoch. This was determined using a statistical test to ask whether each neuron’s activity encoded each particular behavioral feature (see Methods). Gray bar indicates significant encoding of the behavioral feature. (G) Example neurons from the recording depicted in panels (A-G), shown alongside NSM activity. The top two neurons (red) have a positive correlation with NSM. The next two neurons (blue) have a negative correlation with NSM. The two neurons below that have positive (bottom left) and negative (bottom right) correlations with NSM’s derivative (direct overlays of these neurons and NSM derivative are in Fig. S7D). Finally, the neuron on bottom (gray) has no relationship with NSM. Inset text describes properties of the relationship of each neuron with NSM. (H) The correlation of NSM with each of the behavioral parameters listed on the x-axis, measured via a Spearman correlation. Each dot depicts the results from a different dataset (seven total recordings). Red dots indicate that the correlation was significant at p<0.05 using the statistical procedure described in the main text (comparing actual correlation coefficient to those obtained from a distribution of shuffled, synthetic controls; also see Methods). (I) An analysis of how neurons that encode different motor programs (during the pre-food-encounter epoch) are associated with NSM activity (in the post-food-encounter epoch). This provides a measure of how NSM activity is associated with altered activity in core behavioral networks. Here, all neurons recorded across seven different recordings are pooled and shown. **p<0.01, chi-squared test shows a significantly different distribution of NSM-associated neurons in forward velocity and head curvature groups, compared to non-encoding group. Feeding is omitted because there were not a sufficient number of neurons detected to warrant meaningful analysis (Note: this is not due to there being no feeding; it is because the variance in feeding behavior during pre-food-encounter was extremely low. Because feeding was at a nearly constant rate over time, the encoding model had no way to distinguish whether a given neuron encoded feeding).

We next examined brain-wide activity in these recordings (an example dataset is shown in Fig. 7D). First, we determined the main modes of neural dynamics across all the recorded neurons after animals encountered the food patch. We again restricted this analysis to the on-food time period to focus on the main modes of dynamics during ongoing serotonin release. We used principal component analysis (PCA) to extract these main modes of dynamics (Fig. 7E). In every dataset, one of the top three PCs (i.e. the axes of neural activity that explain the most variance in activity across the neurons) was strongly correlated with NSM’s activity (Fig. 7E). In addition, in most animals, one of the other top three PCs was a close match to the derivative of NSM’s activity (Fig. 7E). This suggests that the activity of NSM, as well as the derivative of its activity, are associated with a large portion of the variance in neural activity across the brain when animals encounter food and begin feeding.

To characterize these widespread changes in brain activity more precisely, we examined how the activity of each neuron in each recording was related to NSM activity (example dataset in Fig. 7F; data summary in Fig. 7I). We again restricted our analysis to the post-food-encounter time epoch. The relationship between NSM activity and the activity of any other neuron could be temporally precise (i.e. an immediate time-locked association) or could be temporally lagged. Therefore, we used the following approach to examine whether a given neuron had activity dynamics associated with NSM. We convolved NSM’s activity with a set of temporal filters that could result in, at one extreme, NSM’s original activity and, at the other extreme, a smoothened and shifted version of NSM’s activity (-/+ 10sec). For each neuron simultaneously recorded with NSM, we asked which of these filtered NSM traces was most strongly correlated with it and used an analysis of shuffle controls to test for significance (see Methods). Given that NSM’s derivative was a prominent mode of brain dynamics, we also performed a parallel analysis where NSM’s activity was convolved with filters that take its derivative (NSM activity was not correlated with its own derivative, Fig. S7B-C). Across animals, 33% of the neurons that we recorded showed a significant association with NSM and 21% showed a significant association with its derivative (45 ± 9.1% of neurons were associated with NSM in at least one of these manners). Neurons displayed positive or negative correlations with NSM, though negative correlations were far more prevalent. The optimal kernels to explain the temporal relationships between NSM and other neurons were in some cases quite different: some neurons had immediate time-locked associations with NSM, while others lagged NSM activity (examples in Fig. 7G; see also Fig. S7D for more information about the neurons associated with NSM derivative). Overall, these results show that a large fraction of neurons across the *C. elegans* brain display activity patterns associated with NSM activity. The relationships between NSM and these neurons can be quite different: excitatory, inhibitory, time-lagged, or related to the derivative of NSM activity.

We examined this set of NSM-associated neurons more closely. Specifically, we asked which behavioral networks they participate in by examining how their activity encoded behavior prior to NSM activation (i.e. during the pre-food-encounter time epoch; Fig. 7F, bottom). To quantify this, we used our recently described modeling approach that can reveal whether a given recorded neuron displays significant encoding of the animal’s velocity, head curvature, or feeding (Atanas et al., 2022). This analysis showed that neurons that encoded forward velocity or head curvature prior to food encounter were significantly more likely to display activity that was inversely correlated with NSM after food encounter (Fig. 7I), suggesting that increased NSM activity was associated with inhibition of the networks that encode locomotion and head bending.

### Mapping NSM-associated brain dynamics onto the defined neuron classes of the *C. elegans* connectome, and relating serotonin receptor expression to functional dynamics

Finally, we mapped out which exact neurons in the connectome showed functional associations with NSM. To do so, we performed additional freely-moving brain-wide recordings using the same microscopy platform. However, we used a strain expressing both the pan-neuronal GCaMP7f transgene and the multi-spectral NeuroPAL transgene described above (we used *otIs670*, a low brightness integrant previously shown to be phenotypically wild type in several respects). After the freely-moving recordings, we immobilized each animal and captured multi-spectral fluorescence. We then used the NeuroPAL images to determine the ground-truth identities of the imaged neurons and registered those images back to the freely-moving data. This method, which we have described in detail recently (Atanas et al., 2022), allows us to determine the identities of neurons recorded during freely-moving GCaMP imaging.

The results from the NeuroPAL recordings were consistent with our other brain-wide recordings. NSM activity was significantly correlated with reduced speed and head movements and a similar fraction of neurons was significantly associated with NSM (29% vs 33% above; NSM derivative-associated neurons were slightly less prevalent, 9% vs 21% above). We examined which neuron classes showed reliable associations with NSM activity across recordings (Fig. 8A shows data pooled from six animals; Fig. S8A shows all data from individual animals). When pooled together, sensory-, inter-, and motor-neurons did not differ in their association with NSM (Fig. 8B). Some sensory neurons showed a positive correlation with NSM (AWB, BAG) while others showed a negative correlation (AWC, OLQ). This suggests that ongoing changes in NSM activity on food are associated with bi-directional changes in activity across different sensory channels. Pharyngeal neurons were significantly more likely to be positively correlated with NSM (Fig. 8B), consistent with the positive relationship between NSM and the feeding behavior controlled by the pharyngeal system. We also analyzed whether neurons that displayed activity functionally coupled to different behaviors showed differential coupling to NSM. Matching the above results, locomotion- and head bending-encoding neurons were more likely to be negatively correlated with NSM (Fig. 8C; encoding properties of neuron classes based on Atanas et al., 2022). In addition, feeding-encoding neurons were positively correlated with NSM (Fig. 8C). Thus, NSM-associated brain dynamics map onto functionally defined groups of neurons in the connectome in different ways.

**Figure 8.**
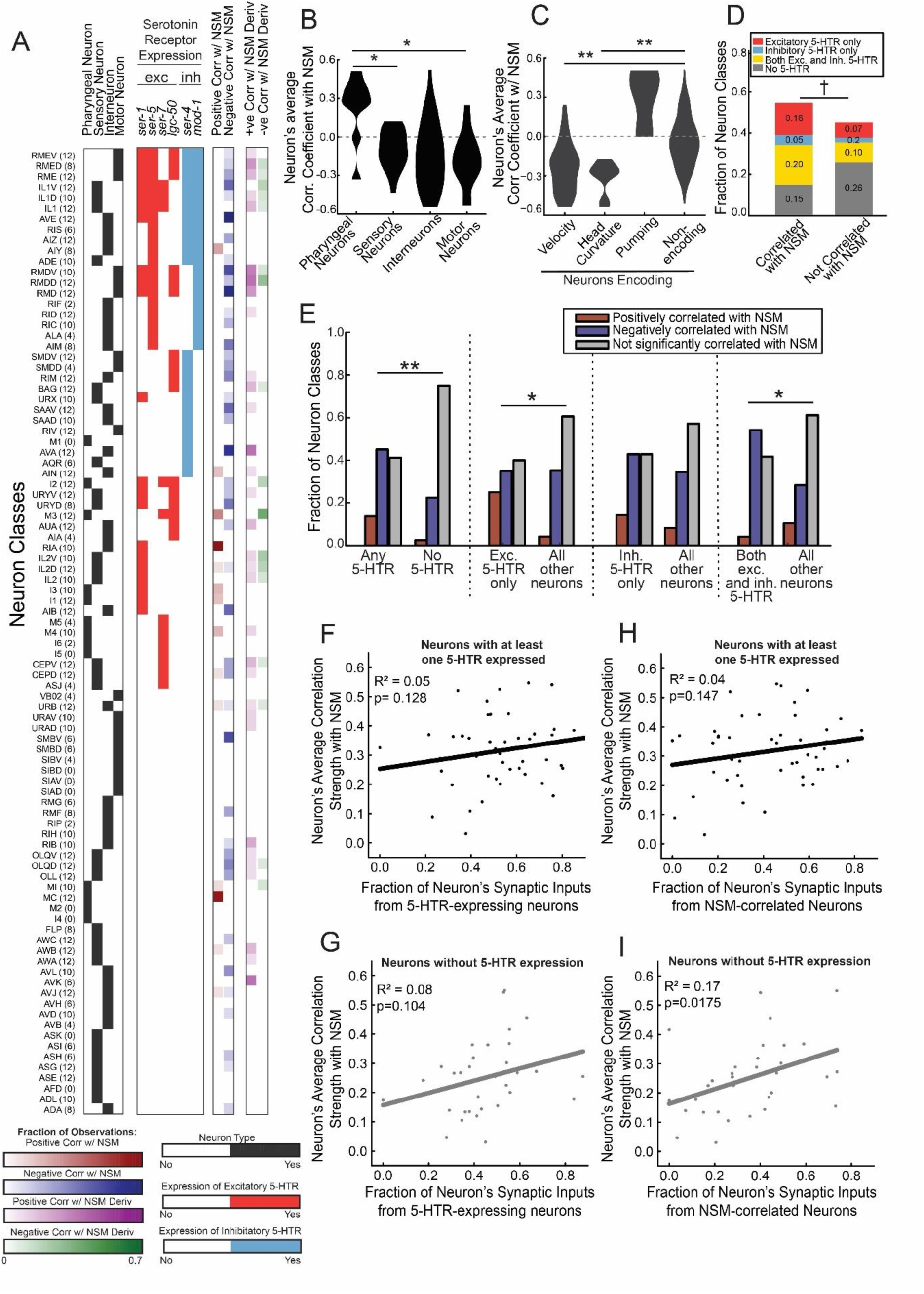
Many neuron classes across the *C. elegans* brain are functionally coupled to NSM, predicted by their serotonin receptor expression patterns, connectivity, and behavioral roles. (A) A list of *C. elegans* neuron classes and how the activity of each is associated with NSM activity. Parentheses indicate the number of recordings analyzed for each neuron (left and right neurons recorded in a single animal are counted here as separate observations). For each *C. elegans* neuron class, we list the following information: (1) Columns 1-4: Whether the neuron is categorized as pharyngeal, sensory, interneuron, or motor neuron. (2) Columns 5-10: The serotonin receptor expression pattern of the neuron, separately highlighting the receptors that are excitatory (red) and inhibitory (blue). (3) Columns 11-12: The fraction of times that the recorded neuron displayed activity positively or negatively correlated with that of NSM (using the statistical procedure described in Main Text and Methods). (4) Columns 13-14: The fraction of times that the recorded neuron displayed activity positively or negatively correlated with that of the derivative of NSM (using the statistical procedure described in Main Text and Methods). (B) Average correlation of the different *C. elegans* neuron classes with NSM, split by the category of neuron. Categorization matches columns 1-4 in panel (A). *, p<0.05, pharyngeal neurons had significantly different correlation with NSM compared to other groups, Brunner-Munzel test (aka Generalized Wilcoxon test). Data distributions are shown as violin plots. (C) Average correlation of neurons to NSM, separated based on how the neuron classes encode behavior. This categorization utilizes data from Atanas et al. (2022). The neurons that were determined to encode each behavior were classified into such groups based on encoding that behavior in >50% of the recordings in Atanas et al. (2022). **p<0.01, neuron groups have different correlations with NSM, Brunner-Munzel test (aka Generalized Wilcoxon test). Data distributions are shown as violin plots. (D) The relationship between a neuron’s serotonin receptor expression profile and whether it showed a significant association with NSM activity. The left bar pools all neuron classes with significant correlations with NSM in >10% of observed recordings (this result was not sensitive to the >10% threshold; same observations were true if threshold was set to >15%); the right bar pools all other neurons. Colored, stacked bars show which neurons expressed which serotonin receptors. †, p=0.0536, chi-squared test showing that relative distributions of neuron types in left and right bars are significantly different. (E) Four separate analyses relating a neuron’s serotonin receptor expression profile to whether it showed a significant correlation with NSM. The four analyses are as follows (from left to right): (1) Comparing neurons with at least one serotonin receptor expressed to all other neurons. (compare first three bars to next three bars) (2) Comparing neurons with only excitatory serotonin receptors expressed to all other neurons. (3) Comparing neurons with only inhibitory serotonin receptors expressed (a very small group of neurons, thus little statistical power) to all other neurons. (4) Comparing neurons with both excitatory and inhibitory receptors expressed to all other neurons. For each analysis, neurons were categorized into three groups, depicted as different color bars. A neuron class was considered to have a significant positive or negative correlation with NSM if a significant correlation was found in >10% of observed recordings. This was not sensitive to exact threshold set, as same observations were true if threshold was set to >15%. *p<0.05, **p<0.01, Chi-squared test showing that relative distribution into the three groups is different between the indicated sets of neurons. (F-G) Relationship between the serotonin receptor expression profiles of each neuron classes’ presynaptic partners and whether that neuron class shows a significant association with NSM. Dots are individual neuron classes. X-axis shows what fraction of the neuron class’ synaptic inputs come from other neurons that express at least one serotonin receptor. Y-axis shows the neuron class’ average correlation with NSM (absolute value of the correlation, since there is no hypothesis with regards to sign of the correlation). The top plot in black (F) shows this analysis for neurons that themselves express serotonin receptors, and the bottom plot in gray (G) shows this analysis for neurons that themselves do not express any serotonin receptors. R-squared and p-values of correlations are shown as insets. (H-I) Relationship between the activity profiles of each neuron classes’ synaptic inputs and whether that neuron class shows a significant association with NSM. Dots are individual neuron classes. X-axis shows what fraction of the neuron class’ synaptic inputs come from neurons that show a significant correlation with NSM. Y-axis shows the neuron class’ average correlation with NSM (absolute value of the correlation, since there is no hypothesis with regards to sign of the correlation). The top plot in black (H) shows this analysis for neurons that themselves express serotonin receptors, and the bottom plot in gray (I) shows this analysis for neurons that themselves do not express any serotonin receptors. R-squared and p-values of correlations are shown as insets.

Taking advantage of these datasets, we examined whether the serotonin receptor expression profiles of neurons could predict their functional correlations with NSM. Indeed, neuron classes that were significantly associated with NSM were more likely to express at least one serotonin receptor (Fig. 8D; Fig. 8E, left). However, a considerable number of neurons did not follow this rule: some neurons with no serotonin receptors were correlated to NSM and some neurons that express serotonin receptors showed no significant correlation (Fig. 8A). We also examined more detailed relationships. For example, neurons expressing only excitatory serotonin receptors were more likely to be positively correlated with NSM (Fig. 8E). Neurons with inhibitory and excitatory receptors were more likely to be negatively correlated with NSM (Fig. 8E). Overall, these results suggest that the neurons with serotonin receptors are more likely to show functional associations with NSM. However, functional correlations with NSM cannot be fully predicted by a neuron’s serotonin receptor expression profile.

We also asked whether information about the presynaptic inputs onto a neuron could predict whether a neuron class was associated with NSM. Neurons with a higher fraction of synaptic inputs coming from neurons that express serotonin receptors were not significantly more likely to be associated with NSM than other neurons (Fig. 8F-G). However, neurons were significantly more likely to be associated to NSM when they had a higher fraction of synaptic inputs coming from presynaptic partner neurons that were functionally associated with NSM (Fig. 8H-I). This association was only significant for the set of neurons that did not themselves express any serotonin receptors (Fig. 8I). This suggests that NSM-associated dynamics in neurons that do not express serotonin receptors are inherited from their presynaptic inputs. Taken together with the above analyses, these results indicate that NSM activity dynamics are associated with widespread changes in brain activity that impact distinct behavioral circuits in different ways, inhibiting the circuits that control movement and activating those that control feeding. The serotonin receptor expression profiles of neurons within these circuits provide a general prediction of how the circuit’s activity will relate to NSM, but additional synaptic interactions in the network further influence the overall activity patterns to give rise to the widespread brain dynamics associated with NSM activity.

## DISCUSSION

Serotonin signaling is critical for the control of behavior and cognition, but our mechanistic understanding of how serotonin acts through diverse receptor types to alter brain activity and behavior remains limited. Here, we examined this problem at brain-wide scale in the *C. elegans* nervous system. We used a behavioral paradigm in which we activated NSM, a feeding-responsive neuron whose extra-synaptic release of serotonin drives behavioral changes associated with foraging, including slow movement and increased feeding. A comprehensive genetic analysis revealed how the full set of serotonin receptors in this organism mediate the effects of serotonin on behavior: one group of “driver” receptors induced slow locomotion upon serotonin release and another group of receptors further modulated these behavioral changes. Surprisingly, we found that different driver receptors supported different dynamical changes in behavior. SER-4 drove transient changes in locomotion in response to abrupt increases in serotonin release and MOD-1 drove sustained slowing during persistent serotonin release. Cell-specific genetic perturbations showed that the sites of expression of the distinct receptors played a key role in conferring their functional properties, and we mapped out all sites of expression of the six receptors across the *C. elegans* nervous system. We performed brain-wide calcium imaging in freely-moving animals with knowledge of cellular identity during serotonin release, providing, for the first time, a view of how serotonin release is associated with changes in activity across the defined cell types of an animal’s brain. This revealed that the sites of serotonin receptor expression can partially predict how individual neurons change activity during serotonin release, but the pervasive brain-wide activity changes that accompany serotonin release extend far beyond these cells to others in the extensive networks that control behavior. Overall, these results provide a global view of how serotonin acts on a diverse set of receptors distributed across a connectome to modulate brain-wide activity and behavior.

We found that NSM serotonin release alters locomotion in a manner that depends on all six known serotonin receptors in the *C. elegans* nervous system. This allowed us to examine the functional logic of how the receptors interact to control behavior. By examining behavioral responses to serotonin release in a panel of mutant animals lacking every possible combination of serotonin receptors, a clear logic emerged. Three serotonin receptors (SER-4, MOD-1, and LGC-50) drive the serotonin-induced locomotion change and three other receptors (SER-1, SER-5, and SER-7) further modulate these effects. The driver receptors are a mixture of ionotropic and metabotropic receptors, and the modulatory receptors are all metabotropic. We identified specific patterns of redundancy and antagonism between combinations of these receptors, which were captured by a model that accurately predicts serotonin-induced slowing based on which receptors are present in a given genotype. Understanding how a given serotonin receptor contributes to a behavioral output and how it interacts with the others will be critical to eventually target serotonin receptors in a rational way for therapeutics. Future studies of receptor interactions in this *C. elegans* model should reveal new underlying mechanisms that give rise to the unique functional roles and interactions of serotonin receptors.

One striking example of a functional difference between the receptors was the distinct behavioral dynamics supported by the MOD-1 and SER-4 receptors. Whereas *mod-1* expression was sufficient for animals to display sustained slowing during persistent NSM activation, *ser-4* expression was only sufficient for animals to transiently slow down in response to abrupt NSM activation. MOD-1 is an ionotropic serotonin-gated chloride channel and SER-4 is a Gi/o-coupled metabotropic receptor (Olde and McCombie, 1997; Ranganathan et al., 2000). Although metabotropic receptors are classically thought to underlie sustained changes in cellular function, in this case the ionotropic receptor supports the sustained behavioral changes. Through receptor swap experiments, we found that these distinct functions are due to the sites of expression of the receptors: when *mod-1* is expressed in *ser-4*-expressing neurons, it only supports transient slowing in response to increased NSM activity, and vice versa. The functional difference between the sets of neurons expressing these two receptors is currently unclear, though it may involve their levels of recurrent excitatory or inhibitory feedback that may further shape their dynamics. Indeed, previous work has suggested that neuromodulation and recurrent circuitry may interact to control stable behavioral states (Kennedy et al., 2020). The finding that there are separable mechanisms that drive behavioral responses to abrupt changes in NSM activity and ongoing tonic NSM activity could have important ethological significance. Sudden increases in NSM activity occur when animals suddenly encounter a better food source and may signal a need to slow down even if there is already high tonic NSM activity (Fig. 1; Iwanir et al., 2016; Rhoades et al., 2019). Continued tonic activity may allow animals to tune their locomotion to smaller ongoing environmental changes. Having both modes of responding may allow animals to maintain sensitivity to different environmental features.

To examine the impact of serotonin on brain dynamics, we recorded native brain-wide activity while animals encountered and ate bacterial food. This is a natural context where NSM is activated to drive slow locomotion. We observed widespread brain dynamics across ∼45% of the brain that was directly associated with time-varying changes in NSM activity or the derivative of its activity. While it is not clear that these effects all reflect the causal influence of NSM on downstream neurons, such widespread effects could be explained by NSM’s extra-synaptic release of serotonin onto the main neuropil of the worm’s brain. Mapping these results onto the connectome revealed changes in brain dynamics that provide a clear intuitive match to NSM-induced behavioral changes. Neurons in the networks that control locomotion and head curvature were inversely correlated with NSM activity, while neurons in the feeding network were positively correlated. Thus, NSM activation is associated with an inhibition of the networks for locomotion and head movement, and increased activity in the feeding network. The patterns of serotonin receptor expression in individual neurons provided a greater than chance prediction of how neurons correlate to NSM, but this was not fully predictive: neurons with no serotonin receptors could still show activity strongly coupled to NSM, and neurons with prominent serotonin receptor expression could show no association. Interestingly, the dynamics of the neurons with no serotonin receptors could be partially predicted by knowledge of how many of their presynaptic partners were associated with NSM. This suggests that the serotonin receptors play an important role in establishing NSM-associated dynamics across the brain, but additional synaptic interactions further shape these patterns.

Collectively, our observations suggest that even finer levels of resolution may be necessary to fully understand serotonergic modulation of brain-wide activity and behavior. For example, we observed a surprising level of co-expression of excitatory and inhibitory serotonin receptors in individual neurons. Two neuron classes even expressed five different serotonin receptor types. One possible explanation for this is that the receptors may localize to different cellular compartments, including different pre- and post-synaptic sites. According to this hypothesis, different synapses along a single neuron could be modulated differently in response to serotonin release. Furthermore, our observations suggest that the brain-wide activity changes that accompany serotonin release are not fully explained by the serotonin receptor expression profiles of the neurons. The precise sites of modulation of each receptor, as well as the way that this modulation interacts with ongoing classical transmission in these circuits, likely gives rise to each serotonin receptor’s unique functional role in modulating brain-wide activity and behavior.

## ACKNOWLEDGMENTS

We thank Matthew Lovett-Barron, Qiang Liu, Cori Bargmann, and members of the Flavell lab for their comments on the manuscript. We thank the Caenorhabditis Genetics Center (supported by P40 OD010440), S. Mitani, the National BioResource Project (NBRP), and Oliver Hobert for sharing strains. E.T. acknowledges funding from the Government of Canada’s New Frontiers in Research Fund (NFRF), NFRFE-2021-00420, and the Natural Sciences and Engineering Research Council of Canada (NSERC), funding reference number RGPIN-2021-02949. S.W.F. acknowledges funding from NIH (GM135413); NSF (Award #1845663); the McKnight Foundation; Alfred P. Sloan Foundation; and the JPB Foundation.

## AUTHOR CONTRIBUTIONS

Conceptualization, U.D., I.N., D.K, and S.W.F. Methodology, U.D., I.N., D.K, J.K., A.A.A, E.B., S.P., Z.W., E.T., and S.W.F. Software, I.N., D.K., J.K., A.A.A., E.B., S.P., E.T., and S.W.F. Formal analysis, U.D., I.N., D.K., and S.W.F. Investigation, U.D., I.N., D.K., M.A.G., J.K., C.E. Writing – Original Draft, U.D., I.N., S.W.F Writing – Review & Editing, U.D., I.N., D.K., and S.W.F. Funding Acquisition, E.T. and S.W.F.

## STAR METHODS

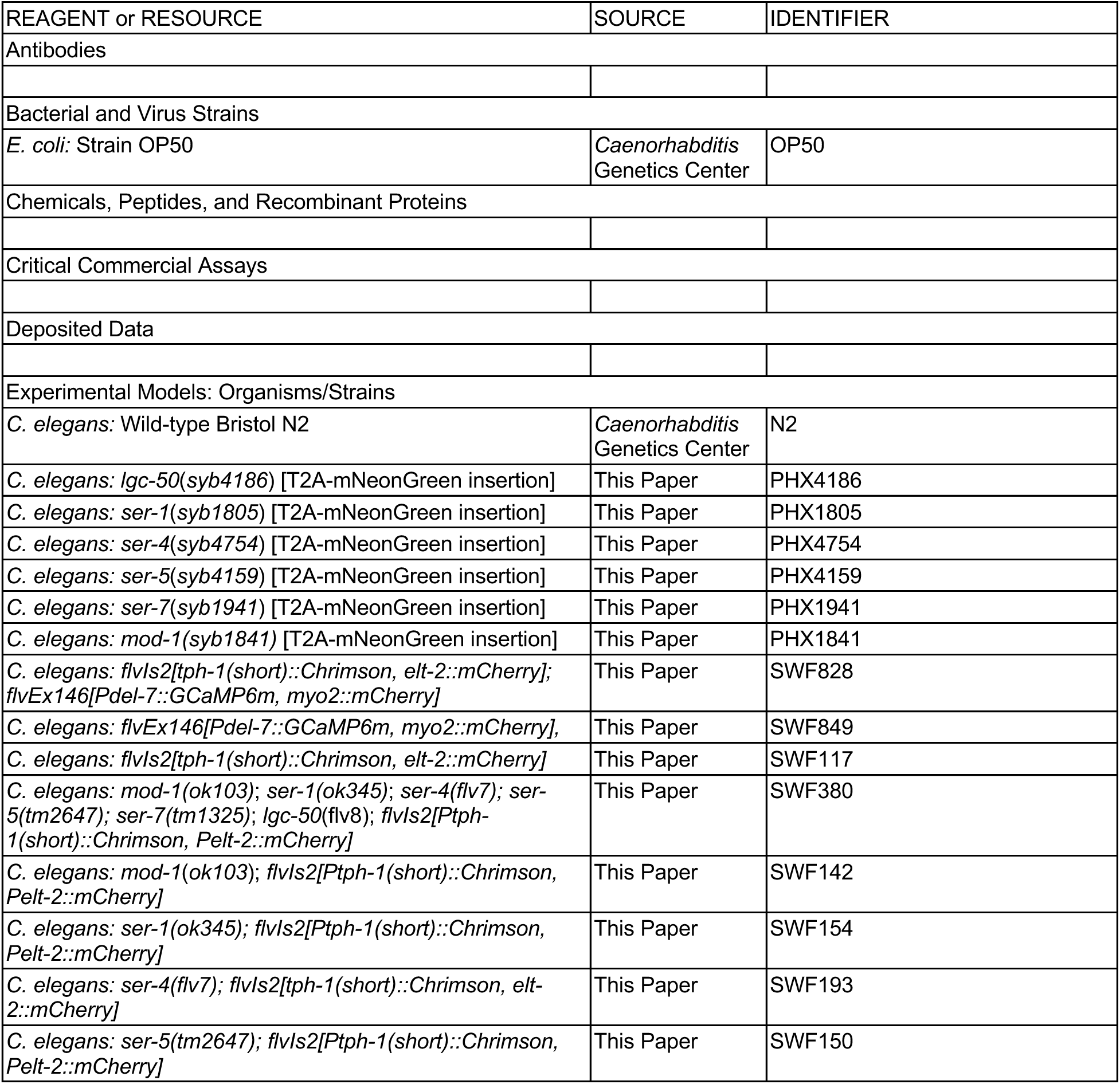
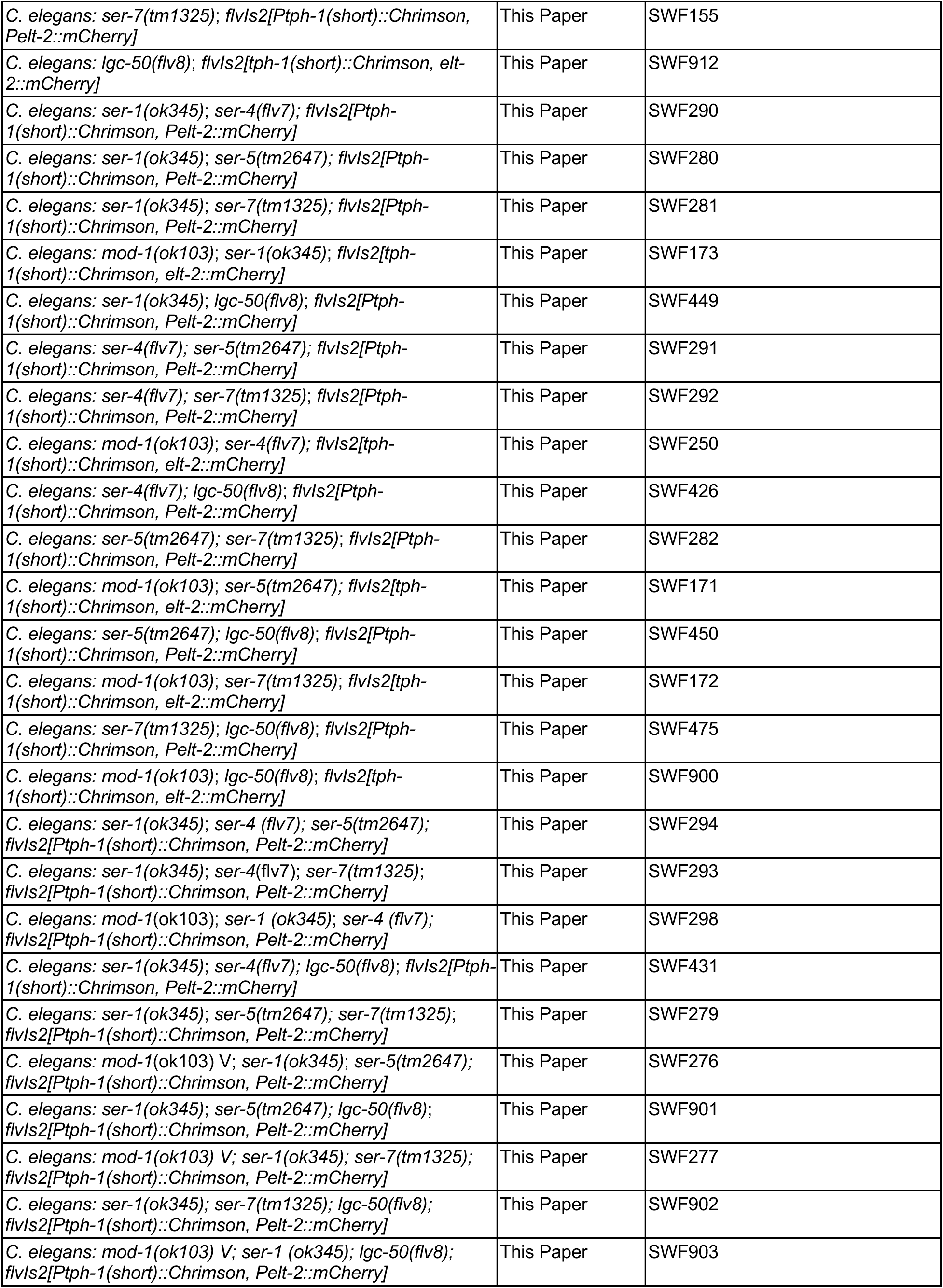
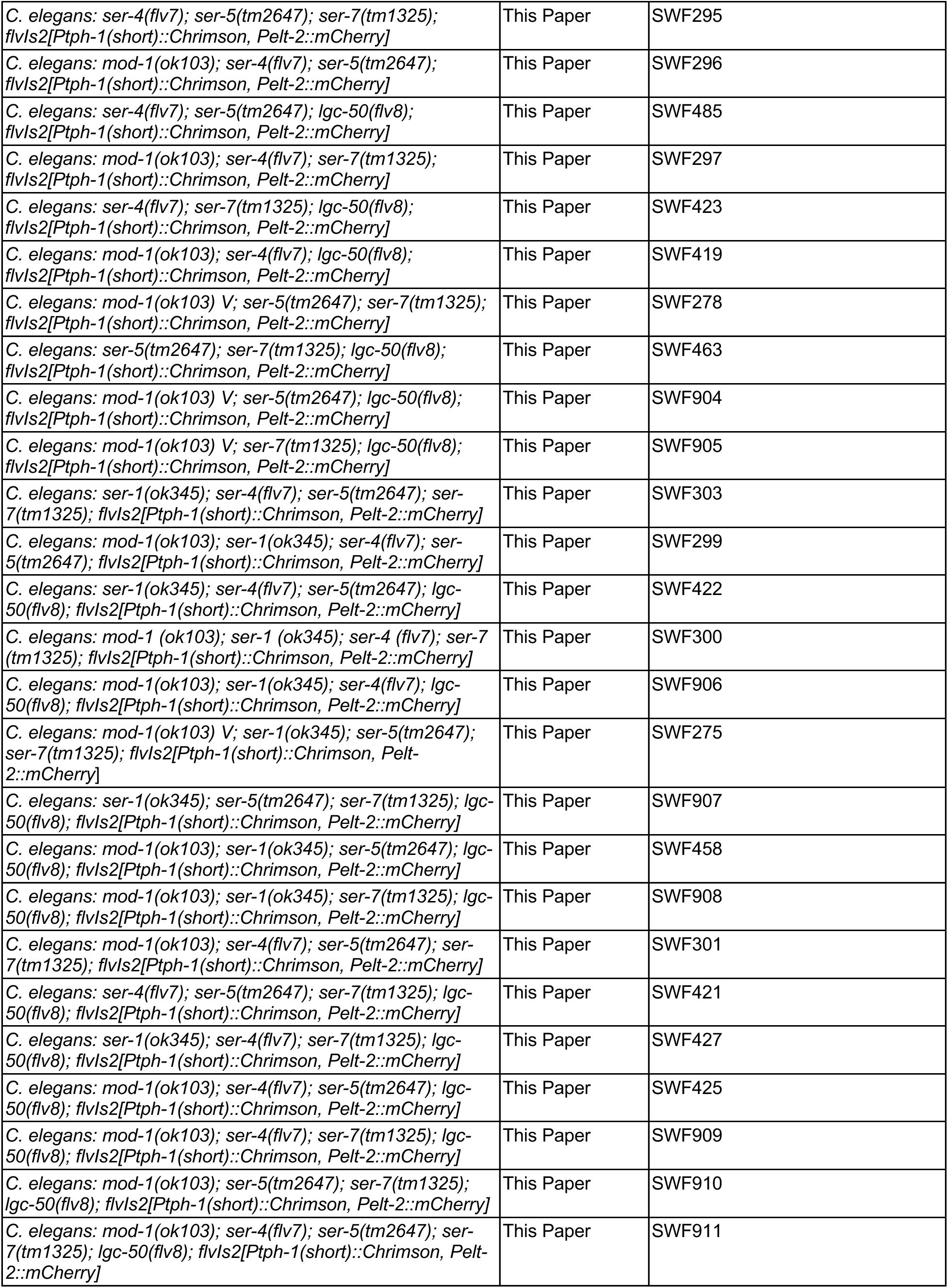
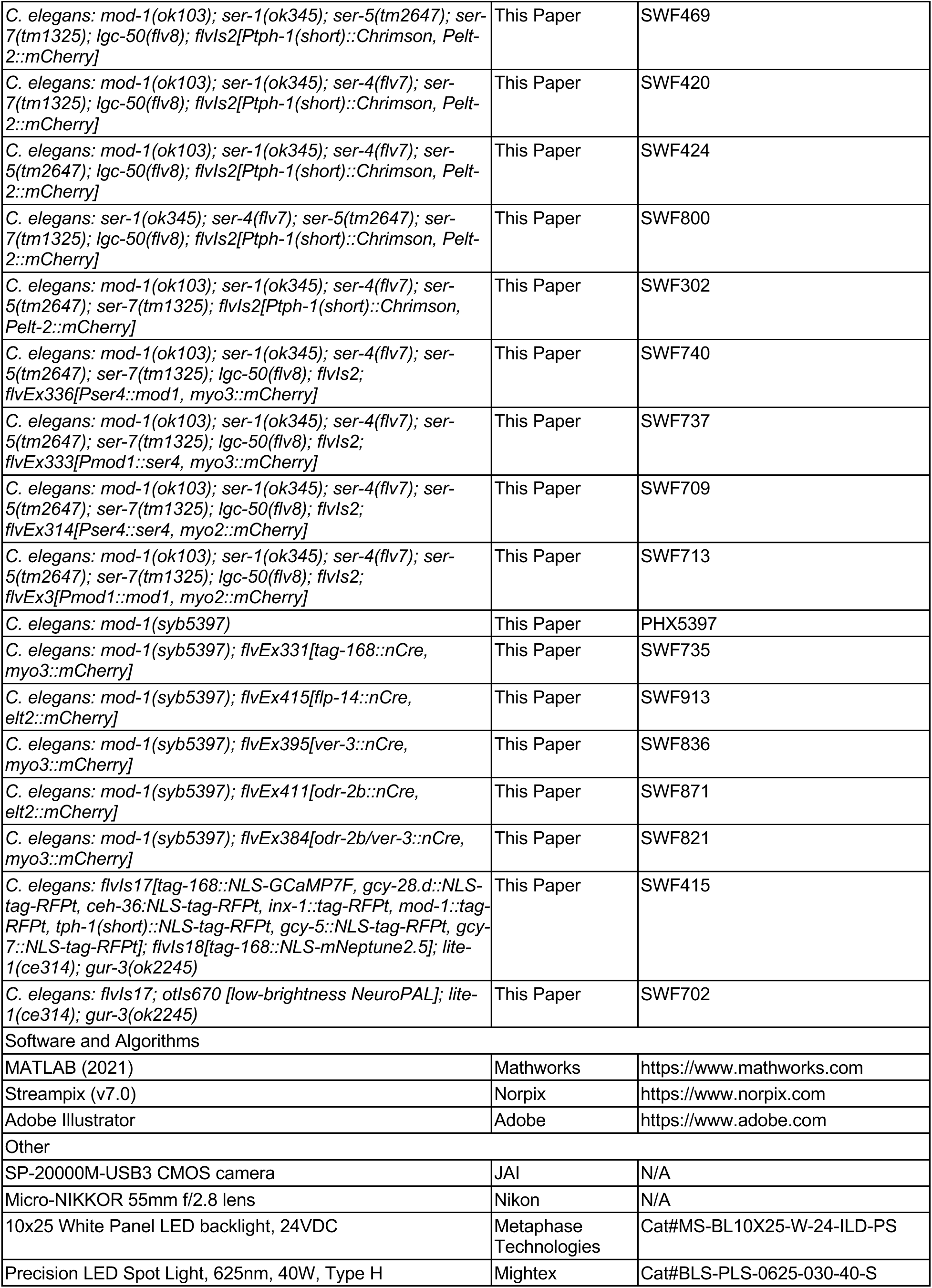
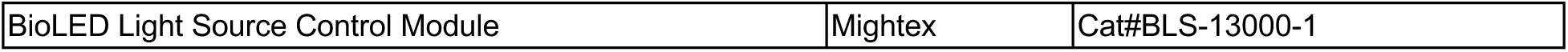
Key Resources Table

### Plasmids

A *tph-1*(short promoter fragment)::Chrimson plasmid that was previously described (Rhoades et al, 2019) was used to drive NSM::Chrimson expression. For transgenic expression of *ser-4* and *mod-1*, the *Y22D7AR.13.1* and *K06C4.6a.1* transcripts were amplified and subcloned into pSM-t2a-GFP. For Cre expression, the following promoters were subcloned into pSM-nCre (which has been previously described; Flavell et al., 2013): *tag-168*, *unc-25, cat-1, nlp-70, ser-2b, tbh-1, ttx-3, flp-14, opt-3, ver-3, odr-2(2b)*.

### New alleles of serotonin receptor genes

We engineered a new predicted null allele of *lgc-50* (*flv8*) using CRISPR/Cas9 gene editing. The mutation changes the sequence TCGAAGTGTTTC***GGC***AAGGATAACACTTG to TCGAAGTGTTTC***ACAGTTTCAGAGAAGTGTTGAG<u>TG</u>***<u>A</u>AGGATAACACTTG (a 3bp deletion and 24bp insertion; bold italics), which introduces an in-frame stop codon (underlined). This mutation is in exon 7, which encodes three of the transmembrane domains of the LGC-50 receptor, suggesting that this allele is likely to be a functional null.

We also engineered a new predicted null allele of *ser-4* (*flv7*) using CRISPR/Cas9 gene editing, because the site of integration for the NSM::Chrimson transgene (*flvIs2*) was closely linked to *ser-4*, such that we were unable to cross existing alleles of *ser-4* with *flvIs2*. The engineered mutation changes the sequence CCTGGAACCTCGG***CG***TCGTGGTCTGTGACTT to CCTGGAACCTCGG***TCTG<u>TGA</u>CT***TCGTGGTCTGTGA (a 2bp deletion and 9bp insertion; bold italics), which introduces an in-frame stop codon (underlined). This mutation is in exon 3 of the gene, which encodes two full transmembrane domains and part of a third, suggesting that this allele is likely to be a functional null.

### Single-neuron calcium imaging of NSM

Single-neuron calcium imaging of NSM was performed as previously described (Rhoades et al., 2019). One day old adult animals expressing GCaMP6m only in the NSM neuron were imaged under a widefield epifluorescence microscope using 10ms exposures at 10 fps. A 4x/0.2NA objective and Andor Zyla 4.2 Plus sCMOS camera were used for imaging. Animals were imaged on flat nematode growth media (NGM) agar pads seeded with *E. coli* OP50 bacteria as described in the main text. Agar imaging pads were prepared immediately before imaging. Neurons were segmented and tracked using custom ImageJ scripts. For experiments with combined NSM::Chrimson stimulation, animals were grown on 5uM all-trans-retinal the night before, imaged at reduced blue light intensity while immobilized with 5mM tetramisole, and red light was delivered via a 617nm red LED light (Mightex).

### NSM-induced locomotion behavioral assays

Locomotion recordings via multi-worm tracker during optogenetic stimulation were conducted using previously described methods (Rhoades et.al, 2019). L4 animals were picked to a 6 cm plate seeded with 200uL OP50 plus 50um ATR 16-24 hours prior to recording. The day of experiments, young adult animals were washed with 750 uL of M9 buffer twice and then placed on 10 cm plates. These 10 cm NGM plates had a thin filter paper ring dipped in 0.02M copper chloride solution. As heavy metals act as an aversive cue to the animals, this served to prevent animals roaming out of the camera’s field of view. For ‘fasted’ condition experiments, animals were washed twice, then placed onto a 6 cm NGM plate with no bacterial lawn present. After 3 hours on this no-food plate, animals were washed twice and recorded immediately on the 10 cm NGM plate with no food.

Streampix software was used to record animals at 3 fps. 625nm light illumination from a Mightex LED was applied at defined times in the video. JAI SP-20000M-USB3 CMOS cameras (41mm, 5120×3840, Mono) with Nikon Micro-NIKKOR 55mm f/2.8 lenses were used. Backlighting was provided by a white panel LED (Metaphase Technologies Inc. White Metastandard 10” X 25,” 24VDC). The recordings were analyzed via custom MATLAB scripts.

### NSM-induced multi-behavioral assays

To record multiple *C. elegans* motor programs, we used a previously described custom microscopy system for single-worm tracking (Cermak et al, 2020). L4 animals were picked onto a 6 cm Nematode Growth Medium (NGM) plate seeded with 200 uL OP50 plus 50 um ATR. Animals fed on this plate in the dark overnight (12-24 hours). The next day, single one-day old adult animals were picked to a 10 cm NGM plate, which had a cut ring of thin film paper, which was soaked in 0.02M Copper Chloride solution. Animals were then placed on a custom tracking microscope that recorded brightfield images of the animal at 20 fps (detailed in Cermak et al., 2020). A 532nm laser provided illumination at defined times in the video. Using custom R Studio scripts, velocity, pumping, head angle, defecation, and egg-laying were calculated for each animal. Custom MATLAB scripts were then used to compile and plot data.

To calculate head oscillations in Fig. 1, head angle data from each animal was analyzed in the spectral domain using the MATLAB Signal Processing Toolbox. To calculate head angle, the animal was divided into 500 equally spaced segments from nose to tail, and head angle was defined as bending around a pivot point at the 39^th^ segment. For this analysis, cubic spline interpolation was used to generate points for small stretches of frames in which angle data was not recorded by the tracker. This was done in order to generate longer stretches of continuous data, which allowed us to achieve better frequency-domain resolution. For each recorded worm, the power spectral density estimate of the worm’s head angle was calculated for each individual laser-off time stretch using the multitaper method. These were then averaged across animals in order to obtain the average power spectral density during baseline behavior. During baseline behavior, we found a peak in power spectral density in the 0.3-0.7 Hz range, corresponding to the frequency of typical oscillatory off-food movement under standard recording conditions. We defined this frequency band as the “roaming band”. We then quantified the average signal power across these frequencies during laser-off and laser-on time stretches to quantify the animals’ head oscillations that accompany high speed movement.

### Food encounter behavioral assays

The food patch encounter behavioral assay was adapted from Iwanir et al. (2016) and Rhoades et al. (2019). Wild-type and mutant animals were picked as L4s to 6mm plates with 200 ul OP50 lawn 16-24 hours prior to experiments. NGM was prepared in 86×128 mm plates (Thermo) and these plates were seeded with OP50 to create a bacterial lawn covering approximately 2/3 of the plate 16 hours before the recordings.

On the day of experiment, worms were gently washed twice with 750 ul M9 buffer to remove any bacterial residues. For the fed conditions, worms were then transferred to the assay plates for recording. For fasted conditions, worms were transferred to a 6mm plate without any OP50 and kept for 3 hours prior to recording.

Streampix software and the hardware described above were used to record animals at 3 fps for 1 hour. Videos were analyzed and data was plotted via custom MATLAB scripts.

### Determining the identities of labeled neurons in the serotonin receptor expression reporter lines

The six T2A-mNeonGreen reporter strains were constructed by inserting a T2A-mNeonGreen coding sequence immediately before the stop codons of *mod-1, ser-1, ser-4, ser-5, ser-7* and *lgc-50* via CRISPR/Cas9 gene editing.

To determine the identities of neurons with serotonin receptor expression, each reporter line was crossed into the NeuroPAL strain. For imaging, one-day old adult animals were picked without any OP50 and immobilized on a flat NGM imaging pad with 100um sodium azide (Sigma Aldrich). Imaging was performed on a Zeiss LSM900 confocal microscope. Imaging parameters and neural identification strategies were conducted as described by Yemini et.al (2021).

### Permutation tests related to receptor expression patterns

To calculate enrichment of serotonin receptors in the Rich Club (Towlson et al., 2013), each stratum (Moyle et al., 2021), and each neuron type, we conducted permutation tests. For a given group of interest 𝐺, (e.g. a specific stratum) of size 𝑁, we randomly selected 𝑁 neurons from the pool of all relevant neurons. For the strata, this pool is the neuropil, for the Rich Club and neuron type it is the non-pharyngeal system. We noted the number of each of the six serotonin receptors in this random sample, 𝑆_!,#_, where 𝑖 is taken from the set {ser-1, ser-4, ser-5, ser-7, mod-1, lgc-50}. We compared the random observation to 𝑆_!,$_, the number of each serotonin receptor in 𝐺. We repeated this random sampling 1000 times. The fraction of random samples for which 𝑆_!,#_ > 𝑆_$_ provides an estimate of the likelihood of observing a value as high (or low) as 𝑆_!,$_.

We calculated the likelihood of observing multiple serotonin receptors in the same single cells via a similar permutation test. For a given neuron containing two different serotonin receptors 𝑖 and 𝑗, we noted the total number of each receptor in the whole network (non-pharyngeal system). We then randomly shuffled the locations of these receptors, such that the same number of receptors were placed across different neurons. We repeated this randomization 1000 times, and counted the number of instances where the serotonin receptors 𝑖 and 𝑗 co-occurred in the same neuron more often than in the real network.

### Linear model for how serotonin receptors interact to drive slow locomotion

We built a linear model to predict the level of NSM-induced slowing across the full panel of serotonin receptor mutants (64 total genotypes). The model with no interaction terms predicted the minimum speed of animals during NSM stimulation using an intercept and six predictor terms, one for each serotonin receptor gene indicating whether it was present (value=1) or absent (value=0) in a given genotype. This model and other versions of it described below were trained using L1-penalized (lasso) maximum likelihood fits, selecting the lambda (regularization) parameter that optimized the cross-validated performance for the particular model. For the linear model with interaction terms, these terms had the form of describing the joint presence of two or more serotonin receptors in a given genotype (for example, an interaction term could describe whether both *ser-4* and *mod-1* were present (value=1) versus if either or both receptors were absent (value=0)). We constructed a model with interaction terms in the following manner. The model initially contained no interaction terms (just the six predictor terms described above). We then asked whether interaction terms describing the joint presence of two receptors (all 2-mer combinations were tested) could be justified based on the data. To do so, we obtained model parameters for a model trained on 3 genotypes: the sextuple mutant and the two quintuple mutants with each of the two receptors (for the 2-mer in question) present. We then used these parameters to predict the level of slowing in the quadruple mutant with both receptors present and compared this to the data from actual animals. If these two values were significantly different (assessed with empirical p-value that the difference was non-zero), then the presence of an interaction term was justified and it was added to the model. If they were not significantly different, no interaction term was added to the model. After all 2-mers were evaluated, they were added to the model and included in a similar analysis to test whether addition of 3-mer interaction terms could be justified (i.e., this evaluation was based on a model that already included the necessary 2-mer interaction terms, and asked whether beyond those any 3-mer interaction terms also needed to be added). This was iterated again for 4-mers. This iterative approach was chosen since it would favor simpler, more parsimonious interaction terms. Once this model was complete, we attempted to discard interaction terms that were not fully necessary in the full model. Specifically, terms that when trained on the full set of genotypes had coefficients that were not significantly different from zero (assessed with empirical p-value) were discarded. This was also to favor the simplest, most parsimonious model.

### Whole-brain calcium imaging recordings

#### Transgenic animals

We recorded data from two transgenic strains that have been previously described. SWF415, which expresses: *flvIs17*: *tag-168::NLS-GCaMP7f*, along with *NLS-TagRFP-T* expressed under the followed promoters: *gcy-28.d, ceh-36, inx-1, mod-1, tph-1(short), gcy-5, gcy-7*; and *flvIs18*: *tag-168::NLS-mNeptune2.5*. And SWF702, which expresses: *flvIs17*: described above; and *otIs670*: low-brightness NeuroPAL (Yemini et al., 2021).

#### Microscope

The microscope used for brain-wide calcium imaging has been previously described. Briefly, the microscope simultaneously images the animal via two light paths, one above and one below the animal. The light path used for all fluorescence imaging is an Andor spinning disk confocal system built on a Nikon ECLIPSE Ti microscope. Laser lines were: 150 mW 488 nm laser, 50 mW 560 nm laser, 100 mW 405 nm laser, or 140 mW 637 nm laser. Excitation light passes through a 5000 rpm Yokogawa CSU-X1 spinning disk unit with a Borealis upgrade (dual-camera). A 40x water immersion objective (CFI APO LWD 40X WI 1.15 NA LAMBDA S, Nikon) with an objective piezo (P-726 PIFOC, Physik Instrumente (PI)) was used to image the volume of the worm’s head. A quad dichroic mirror directed light emitted from the specimen to two separate sCMOS cameras (Andor Zyla 4.2 PLUS sCMOS), which had in-line emission filters (525/50 for GCaMP/GFP, and 610 longpass for mNeptune2.5; NeuroPAL filters described below). Data was collected at 3 × 3 binning in a 322 × 210 region of interest in the center of the field of view, with 80 z planes collected at a spacing of 0.54 um, resulting in a volume rate of 1.7 Hz.

The light path for behavior imaging was in a reflected brightfield (NIR) configuration. Light from a 850-nm LED (M850L3, Thorlabs) was collimated and passed through a 850/10 bandpass filter (FBH850-10, Thorlabs). Illumination light was reflected towards the sample by a half mirror and was focused on the sample through a 10x objective (CFI Plan Fluor 10x, Nikon). Image from the sample passed through the half mirror and was filtered by another 850-nm bandpass filter of the same model. The image is captured by a CMOS camera (BFS-U3-28S5M-C, FLIR).

The microscope utilized a closed-loop tracking system to keep the animal in view. For closed-loop tracking, NIR brightfield images were analyzed at a rate of 40 Hz to determine the location of the worm’s head, identified via a custom-trained network with transfer learning using DeepLabCut (Mathis et al., 2018). This network identified the location of three key points in the worm’s head (nose, metacorpus of pharynx, and grinder of pharynx). The tracking target that was kept centered in the field of view was a point halfway between the metacorpus and grinder (central location of neuronal cell bodies). Additional details are available in the original article to reporter this microscope.

#### Recording procedure

Animals were raised on *E. coli* OP50 and imaged as one day-old adults. Each animal was mounted on a thin, flat agar pad (2.5 cm x 1.8 cm x 0.8 mm) with NGM containing 2% agar. *E. coli* OP50 that had been seeded on agar plates 3 days beforehand was spread on the agar pad with a sterile inoculating loop (Thermo Scientific™), after which 4 uL of M9 buffer was added to the center of the agar pad. The animal was then transferred to the drop of M9 buffer. At each corner of the agar pad, 1 uL of 80 um microbeads suspended in M9 buffer were added as vertical spacers, on top which a #1.5 glass coverslip was added.

Each SWF415 animal was starved for 3-4 hours before the mounting procedure. *E. coli* OP50 was spread along three sides of the agar pad such that the animal is ∼0.6 cm away from each lawn boundary. The glass coverslip was lowered gently from the foodless side in order to avoid contaminating the central foodless zone with *E. coli* OP50. Imaging was performed 3 minutes after the mounting procedure.

Each SWF702 animal was starved for 1.5-2 hours before the mounting procedure. *E. coli* OP50 was spread evenly to form a circular lawn of 0.5 cm diameter, and the animal was mounted at the center of the food lawn. Imaging was performed immediately after the mounting procedure to capture the initial phase of food-induced neural activity.

#### Data processing

Whole-brain GCaMP/mNeptune data were processed into normalized calcium traces using the Automated Neuron Tracking System for Unconstrained Nematodes (ANTSUN) data processing pipeline that we have previously described (Atanas et al., 2022).

Briefly, this software package uses the mNeptune signal to identify the neurons in all frames and to register volumes from different time points to one another. The end result is a Fluorescence (F) measurement for each neuron, which is the ratio of GCaMP7F signal divided by mNeptune2.5 signal for that neuron. Behavioral data were extracted from the NIR videos using previously described methods.

#### Procedure for neuroPAL imaging

NeuroPAL recording procedures were carried out as previously described. Briefly, animals were imaged as described above, but they were subsequently immobilized by cooling, after which multi-spectral information was captured. A closed-loop temperature controller (TEC200C, Thorlabs) with a micro-thermistor (SC30F103A, Amphenol) embedded in the agar was used to keep the agar temperature at the 1 °C set point. We then captured a series of images necessary for NeuroPAL-based neural identification:

(1-3) Spectrally isolated images of mTagBFP2, CyOFP1, and mNeptune2.5. CyOFP1 data was collected with 488nm laser excitation under a 585/40 bandpass filter. mNeptune2.5 was collected with 637nm laser excitation and a 655LP-TRF filter. mTagBFP2 was isolated with 405nm laser excitation and a 447/60 bandpass filter.

(4) An image with TagRFP-T, CyOFP1, and mNeptune2.5 (all of the “red” markers) in one channel, and GCaMP7f in the other channel. This image was used for neuronal segmentation and registration to both the freely moving recording and individually isolated marker images. For this image, we used 488nm and 561nm laser excitation. TagRFP-T, mNeptune2.5, and CyOFP1 were imaged with a 570LP filter and GCaMP6f was isolated using a 525/50 bandpass filter.

All images were recorded for 60 timepoints. We increased the signal to noise ratio for each of the images by first registering all timepoints within a recording to one another and then averaging the transformed images. Finally, we created the composite, 3-dimensional RGB image by setting the mTagBFP2 image to blue, CyOFP1 image to green, and mNeptune2.5 image to red as done by Yemini et al. (2021) and manually adjusting the intensity of each channel to optimally match their manual.

The neuron segmentation U-Net used in the ANTSUN pipeline was run on the “all red” image and we then determined the identities of U-net identified neurons using the NeuroPAL instructions. In some cases, neuronal identities could not be determined with certainty due a number of factors including: unexpectedly dim expression of one or more fluorophores, unexpected expression of a fluorophore in cells not stated to express a given marker, and extra cells in a region expressing similar intensities when no other cells are expected. Rarely, multiple cells were labeled as potential candidates for a given neuron and the most likely candidate (based on position, coloring, and marker intensity) was used for analysis.

Finally, the neural identity labels from the RGB image were mapped back to the GCaMP traces from the freely-moving animal by first registering each fluorophore-isolated image to the image containing all of the red markers. The “all red” image was then registered back to the freely moving recording, permitting mapping of neuronal labels back to GCaMP traces.

### Whole-brain calcium imaging data analysis

#### Principal Component Analysis (PCA)

To determine the main modes of neural dynamics in our recordings, we performed PCA on our neural datasets. To do so, we used the F/F20 GCaMP traces and performed PCA on all traces from a given recording using the post-food-encounter time epoch. Analyzing the full datasets (pre- and post-food-encounter) gave rise to qualitatively similar results, but it was more obvious that the PCs arising from that analysis would be related to NSM, which is activated by feeding. By performing the analysis on only post-food-encounter data, we were able to assess whether ongoing brain dynamics on food were related to NSM activity. To relate the resulting principal components to NSM, we performed the analysis described below for single neurons. However, instead of relating NSM to another’s neuron’s GCaMP signal, we correlated NSM activity to each principal component that explained >2% of the overall variance in neural activity.

#### Statistical procedure to determine how NSM activity is associated with behavior

For each brain-wide recording, we examined whether NSM activity was associated with ongoing behavioral changes. Here, we focused on the relationship between NSM and behavior only in the post-food-encounter time epoch. This is because NSM is activated by feeding and analyzing all data (pre- and post-food-encounter) could potentially lead to the spurious conclusion that NSM is correlated with all feeding-induced behaviors, even when the time-locked association is weak. By focusing on the post-food-encounter-time epoch, we were able to ask whether ongoing changes in NSM activity were precisely associated with ongoing behavioral changes.

We examined NSM’s relationship to five behavioral variables: velocity, speed, head curvature, head movement (absolute value of derivative of head curvature), and feeding (i.e. pharyngeal pumping). For each behavior, we performed the following analysis. First, we determined the Spearman correlation between NSM activity and the behavioral variable. To test whether the observed correlation coefficient exceeded values expected by random chance, we compared this value to a distribution of shuffle controls. This distribution was created by obtaining the correlation coefficients between shuffled, synthetic NSM neurons and the behavioral variable being tested. The idea is that the shuffled, synthetic traces would have the temporal properties of NSM activity (autocorrelation, etc), but would be completely de-synced from the behavior so that they estimate the level of correlation expected by chance. The shuffled, synthetic traces were obtained by building an encoding model that accurately predicts NSM GCaMP based on animal velocity (using the procedure in Atanas et al, 2022). We then used random samples of actual recorded animal velocity (from animals recorded under similar conditions; we ensured velocity samples had the same overall velocity distributions as the recorded animals) to synthesize NSM traces and used these for the shuffle controls. Finally, we then determined the empirical p-value of the actual correlation coefficient based on its rank in the control distribution. We favored this approach over the more standard approach of temporally shifting NSM activity relative to behavior, because NSM has a very long autocorrelation, so to obtain independent shifted versions of NSM, its trace needs to be shifted by >100s. Given the durations of our recordings, this makes it infeasible to obtain a sufficient number of independent, shifted NSMs to perform statistical testing (at least 1000 required; preferably more). To ensure that the statistical test was valid, we performed control analyses where we examined whether actual NSM neurons from one animal were correlated to the behaviors of other animals and found that for all permutations of this test they were not.

#### Statistical procedure to determine how each neuron’s activity is associated with NSM activity

To test whether NSM activity in a given recording was associated with the activity of other simultaneously recorded neurons, we performed the following analysis. First, we only considered neurons for analysis if their signal variation, measured as standard deviation of their F/F20 signal, exceeded a value determined to be the boundary between real dynamics and measurement or motion noise. This value was determined based on an analysis of signal variation in the recordings where GFP was recorded in place of GCaMP. All GFP-expressing neurons fell below this value and only <15% of GCaMP neurons were excluded based on this criterion. These neurons were likely inactive under our recording conditions. Next, we wanted to allow for the possibility that NSM could have slightly lagged relationships to other neurons in the data. Thus, we first convolved the activity of NSM with a set of kernels that filter its activity. The kernels varied in their temporal properties, such that, depending on the kernel, it could weigh NSM activity up to 10sec in the past or future. There were 20 such kernels, and therefore this resulted in 20 filtered NSM traces. Then, for a given neuron of interest, we determined which filtered NSM trace had the strongest correlation with the neuron. Then, to test whether this correlation coefficient was significant, we used the procedure described in the above paragraph (for NSM vs behavior analyses) where we obtained a distribution of shuffle controls. For these controls, each shuffled, synthetic NSM trace was put through the same procedure. The idea is that these traces have the temporal properties of NSM activity, but should be completely de-synced from the neuron being analyzed so that this estimates the level of correlation expected by chance. Importantly, the shuffled, synthetic NSM traces were all convolved with the same filters and the highest possible correlation coefficient was the one added to the control distribution. We then determined the empirical p-value of the actual correlation coefficient based on its rank in the control distribution. Multiple comparison correction was applied across all neurons in each dataset using Benjamini-Hochberg set to FDR<0.1. We also implemented a second requirement for neurons to be considered as significantly associated with NSM. Specifically, we aimed to exclude neurons where the variance explained by its correlation with NSM was not separable from levels of measurement or motion artifact noise. To estimate this, we utilized a set of recordings where NLS-GFP was recorded instead of NLS-GCaMP. For each GFP-expressing neuron, we attempted to explain its activity with NSM GCaMP from another recording and then estimated the variance explained in the GFP signal by NSM. This was a typically small number that clearly reflects noise. We selected a threshold for this number that exceeded all of these control measurements. Then, for each actual neuron simultaneously recorded with NSM that passed the shuffle test, we asked whether the variance explained by its correlation with NSM exceeded this threshold value. If it did not, the neuron was not considered to be significantly associated with NSM.

In parallel, we performed an identical analysis using a series of differentiator kernels that take the derivative of NSM. Four possible kernels were used that took the NSM derivative with different time steps. Otherwise, the procedure was identical to the one described above. For both tests, we ran control analyses to ensure the validity of the tests, in which we asked whether NSM from one animal could be correlated to the neurons from another animal and found that <5% of neurons were significant for these analyses (vs 44.9% in actual analyses). All statistical analyses of the whole-brain imaging data were performed using Julia 1.8.0.

#### Determining the encoding properties of recorded neurons

For many of our recordings, it was of interest to determine how each recorded neuron “encoded” behavior. “Encoded” here is defined as displayed GCaMP dynamics that were associated with the behavior. To perform this analysis, we used a previously-described statistical method (Atanas et al., 2022). A detailed description can be found in the cited study, but the conceptual framework is also described here. Briefly, we attempted to fit each recorded neuron with a probabilistic encoding model that attempts to use behavioral predictor terms (for velocity, movement direction, head curvature, and feeding) to describe the neuron’s activity. The form of the model allows neurons to encode behavior over varying timescales and potentially represent multiple behaviors. We fit the model using the probabilistic computing so that we can determine the full posterior distribution of model parameters that could explain the neural/behavioral data. This allows us to establish our confidence in a given model parameter, for example the parameter that explains how much the neuron’s activity reflects head curvature. Finally, we use these posterior distributions to test for significance. In this study, we asked whether each neuron significantly encoded: forward velocity, reverse velocity, dorsal head curvature, ventral head curvature, pumping (in a positive-correlated manner), and pumping (in a negative-correlated manner). To do so, we evaluated all of the points in the posterior distribution, asking whether that set of model parameters indicates tuning to the behavioral parameter of interest (for example, higher activity during forward versus reverse velocity). We then ask whether the null hypothesis of no significant tuning can be rejected (FDR<0.05), by examining the results from all points in the posterior.

---

**Figure S1, Related to Figure 1.**
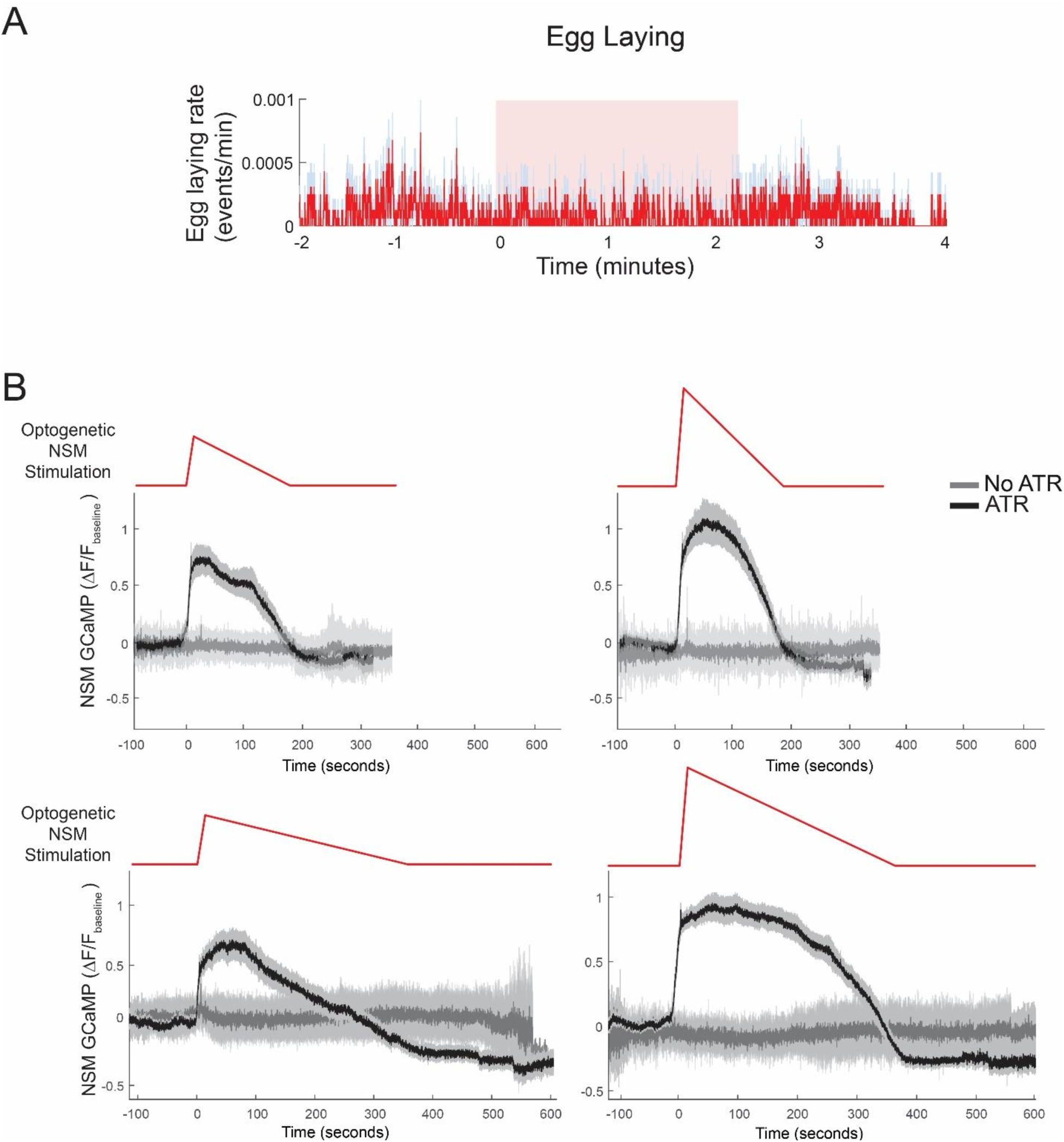
(A) Egg-laying behavior of animals in response to NSM::Chrimson activation. Note that NSM::Chrimson activation has no effect on egg-laying rates. Because of the lack of effect, this was not quantified in the serotonin receptor mutant strains. N=45 stimulation events (3 per animal). (B) NSM GCaMP6m recordings during simultaneous NSM::Chrimson activation. Animals were immobilized with tetramisole and imaged under dim blue light conditions (with short light pulses) to prevent blue light activation of Chrimson. Red light was applied in the indicated temporal patterns to activate NSM::Chrimson. Note that brighter red light led to stronger NSM GCaMP responses and more prolonged red light exposure led to longer NSM GCaMP activity bouts. N=27-59 animals per condition for +ATR conditions; n=10-23 animals for No-ATR conditions.

**Figure S2, Related to Figure 2.**
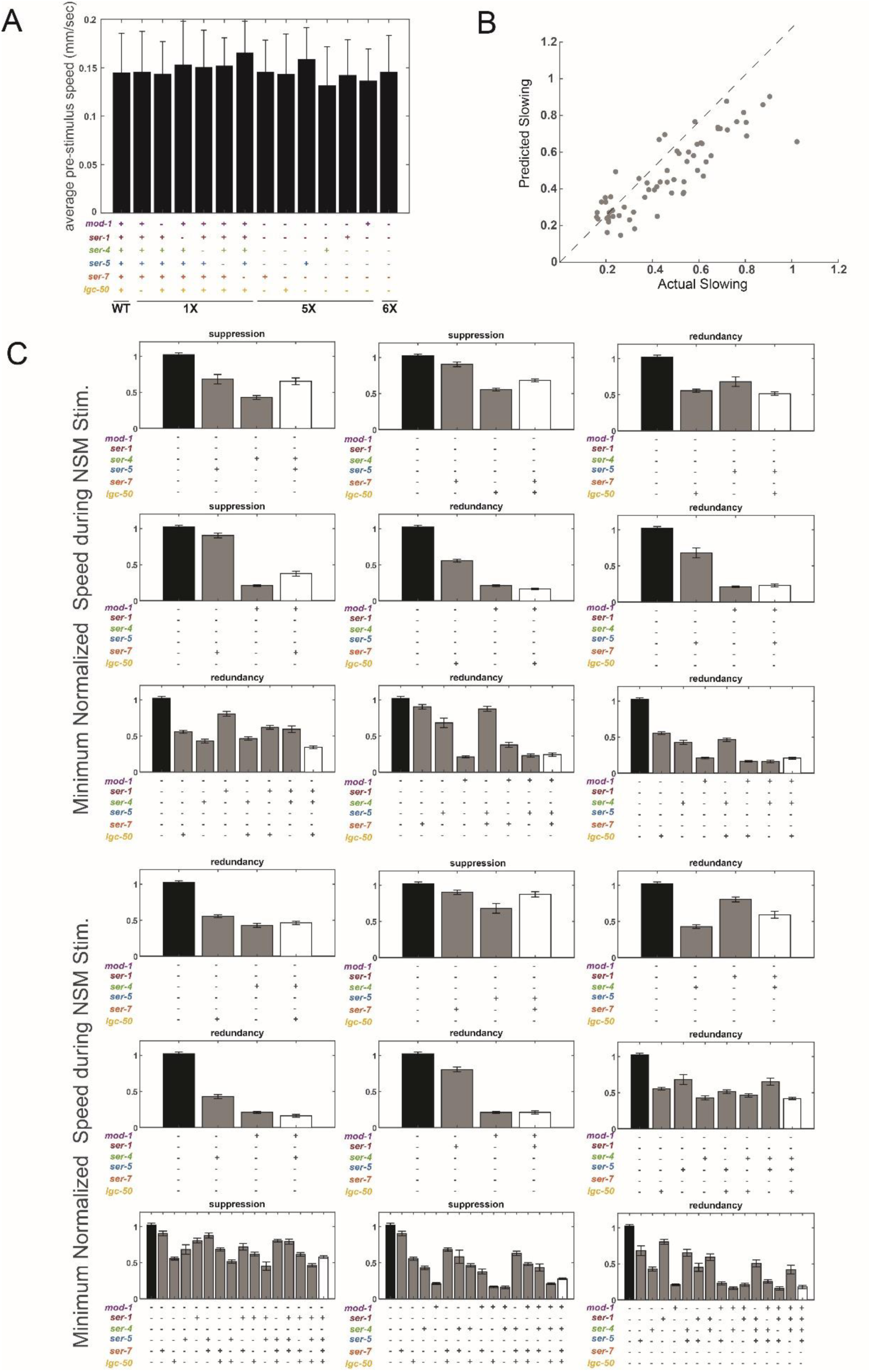
(A) Baseline speeds of animals of the indicated genotypes prior to optogenetic NSM stimulation. Note that baseline speeds are not significantly different from one another. n=45-496 animals per genotype. Data are shown as means ± standard deviation. (B) Performance of the linear model (with interaction terms) that predicts NSM-induced slowing, using a train/test design where the model was trained on data from 63 of the 64 serotonin receptor mutant genotypes and then used to predict the slowing behavior of the withheld genotype. This procedure was repeated 64 times (once for each genotype) and the corresponding dots in the scatter plot are from these trials. (C) Each plot here depicts a set of mutant phenotypes related to a single interaction term that was added to the linear model that predicts NSM-induced slowing behavior based on animals’ genotypes. The statistical criteria to add an interaction term to the model is described in Methods. Here, to make clear what these interaction terms capture about receptor interactions, we display a series of slowing phenotypes in each panel. For each interaction term, we display the NSM-induced slowing behavior of the compound mutant that corresponds to the interaction term. For example, there was an interaction term in the linear model that allows the presence of *ser-5* to inhibit slowing when *ser-4* is present (top left). For this interaction term, we display the slowing behavior of 6X 5-HTR animals, animals with only *ser-4* or *ser-5* present, and animals with both *ser-4* and *ser-5* present. The plot labels indicate the type of interaction between the receptors: suppression or redundancy (no examples of synergistic interactions were detected).

**Figure S3, Related to Figure 3.**
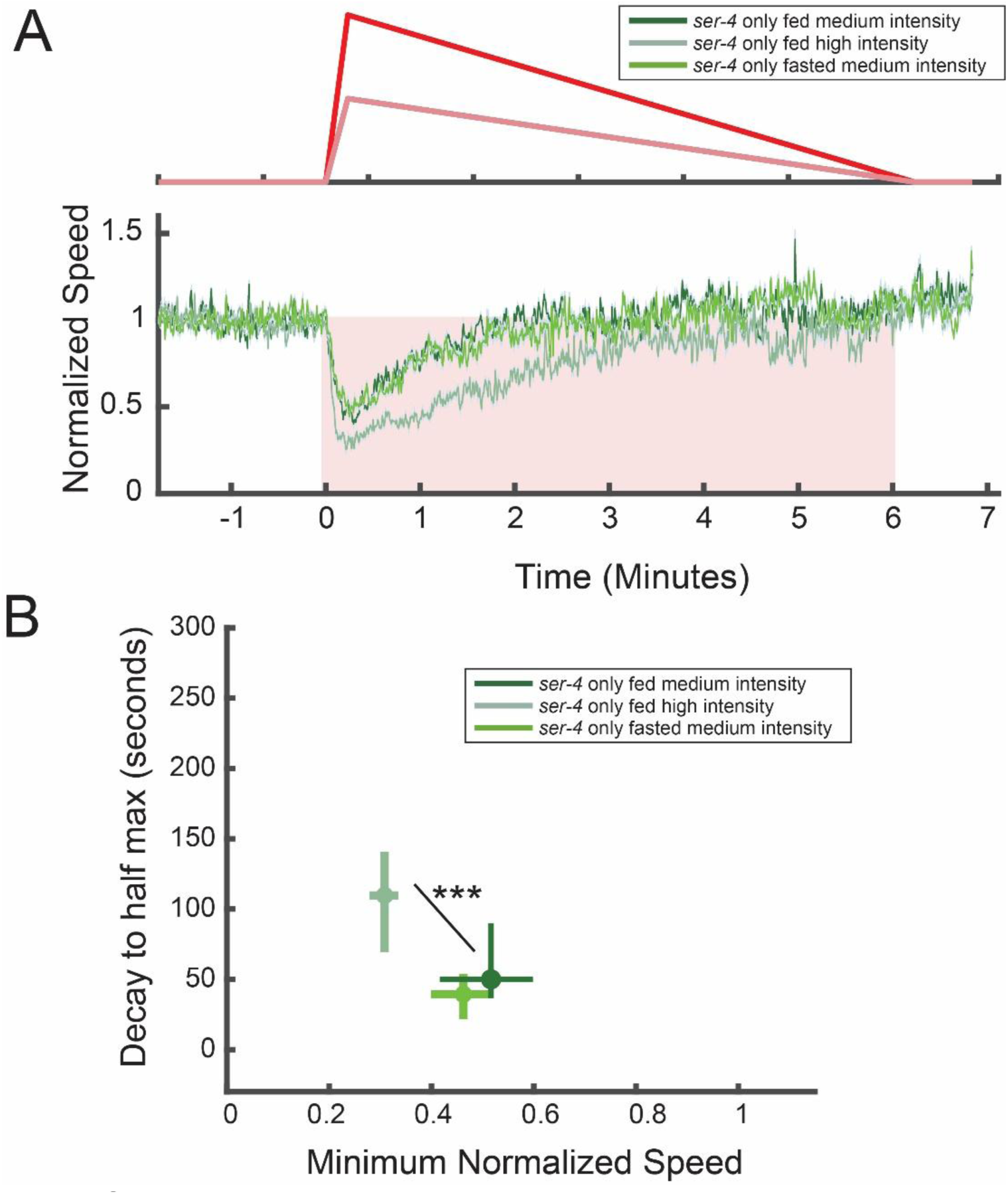
(A) Changes in animal speed for *ser-4*-only mutant animals in response to NSM::Chrimson activation with the indicated waveforms of light. Note that data are from fed or fasted animals, as indicated. Compared to medium-amplitude NSM::Chrimson stimulation in fed animals, fasted animals show no enhancement in slowing. However, stronger-amplitude NSM::Chrimson stimulation in fed animals does lead to a larger change in speed, compared to medium-amplitude stimulation in fed animals. Thus, fasting animals does not exaggerate slowing in *ser-4*-only mutants and this is not due to a saturation effect. Data are quantified in panel (B) (B) Quantification of data in (A). Data are shown as means ± 95% confidence interval. N=83-103 animals per condition. ***p<0.001, empirical p-value that difference between slowing distributions is non-zero.

**Figure S4, Related to Figure 4.**
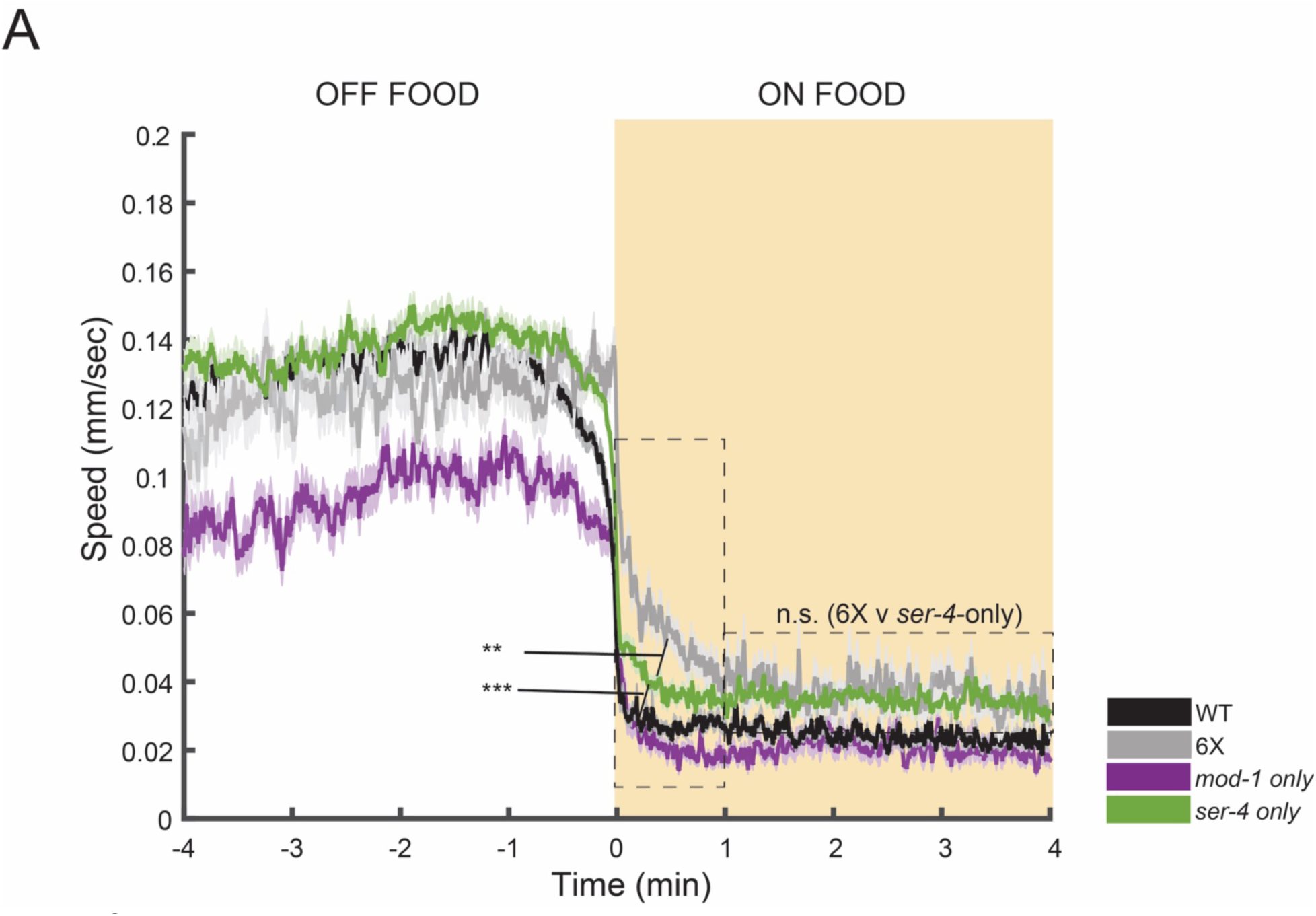
(A) Animal speed for the indicated genotypes shown as event triggered averages surrounding bacterial food patch encounters. Data are shown as means ± SEM. N= 81-221, **p<0.01, ***p<0.001, two-sided Wilcoxon rank sum test between the indicated groups over the indicated time ranges.

**Figure S5, Related to Figure 5.**
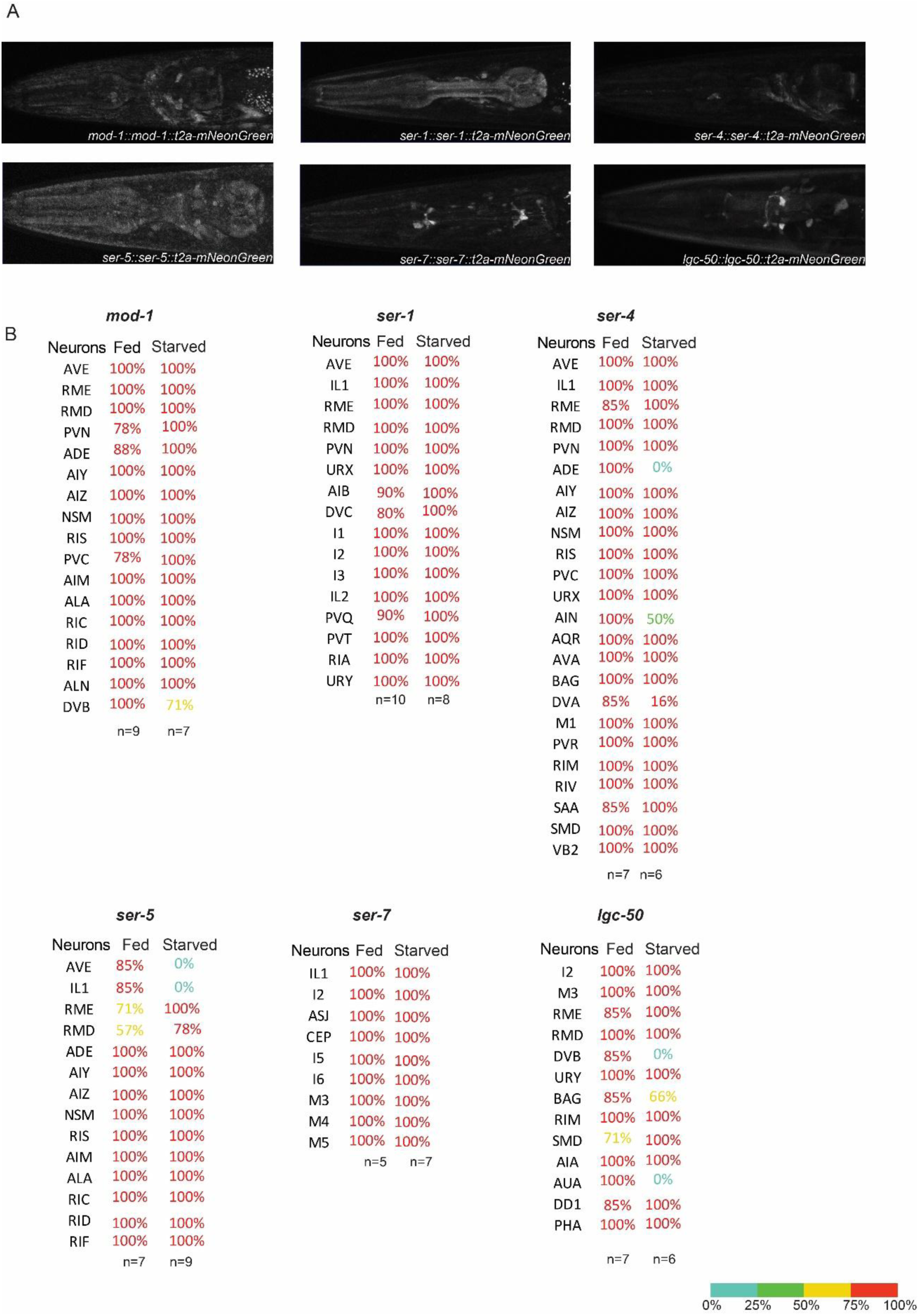
(A) Images of the fluorescent reporters for the six serotonin receptor genes, shown without overlaying NeuroPAL images. Details about construction of the fluorescent reporters are in Fig. 5A. (B) Results of cell identification from the six strains expressing t2a-mNeonGreen reporters from each serotonin receptor gene. For each reporter line, we show the fraction of animals where each mNeonGreen was detected in each cell. Cells that are not listed had 0% detected.

**Figure S6, Related to Figure 6.**
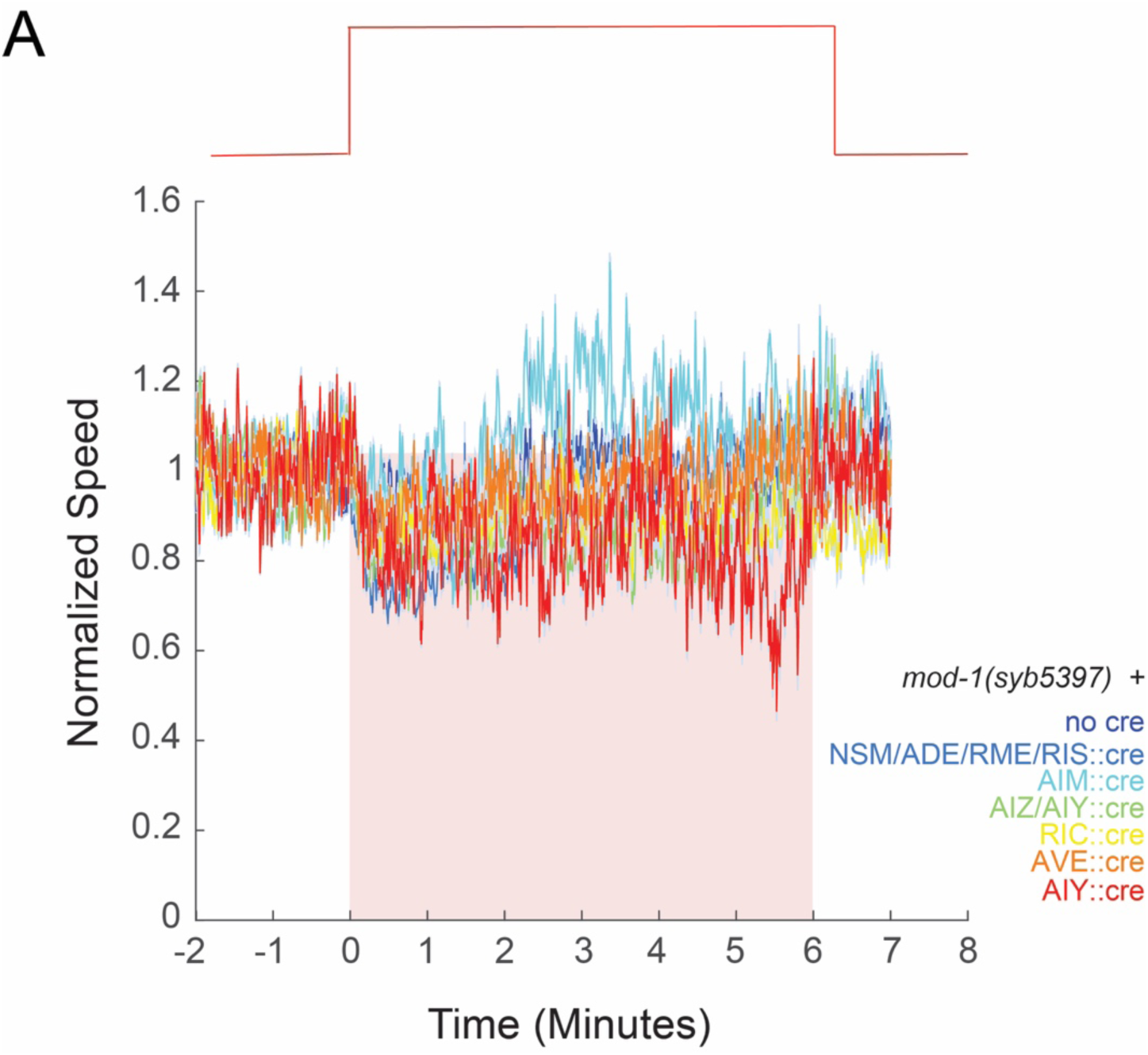
(A) Changes in animal speed in response to NSM::Chrimson stimulation in the indicated *mod-1* rescue strains. Note that genetic rescues in these indicated neurons did not lead to a rescue of NSM-induced slowing, compared to the mod-1 inverted allele. N=29-116 animals per condition. Promoters used for Cre expression were: *Pcat-1* (NSM, ADE), *Punc-25* (RME, RIS), *Pser-2b* (AIY, AIZ), *Ptbh-1* (RIC), *Popt-3* (AVE), *Pnlp-70* (AIM), *Pttx-3* (AIY).

**Figure S7, Related to Figure 7.**
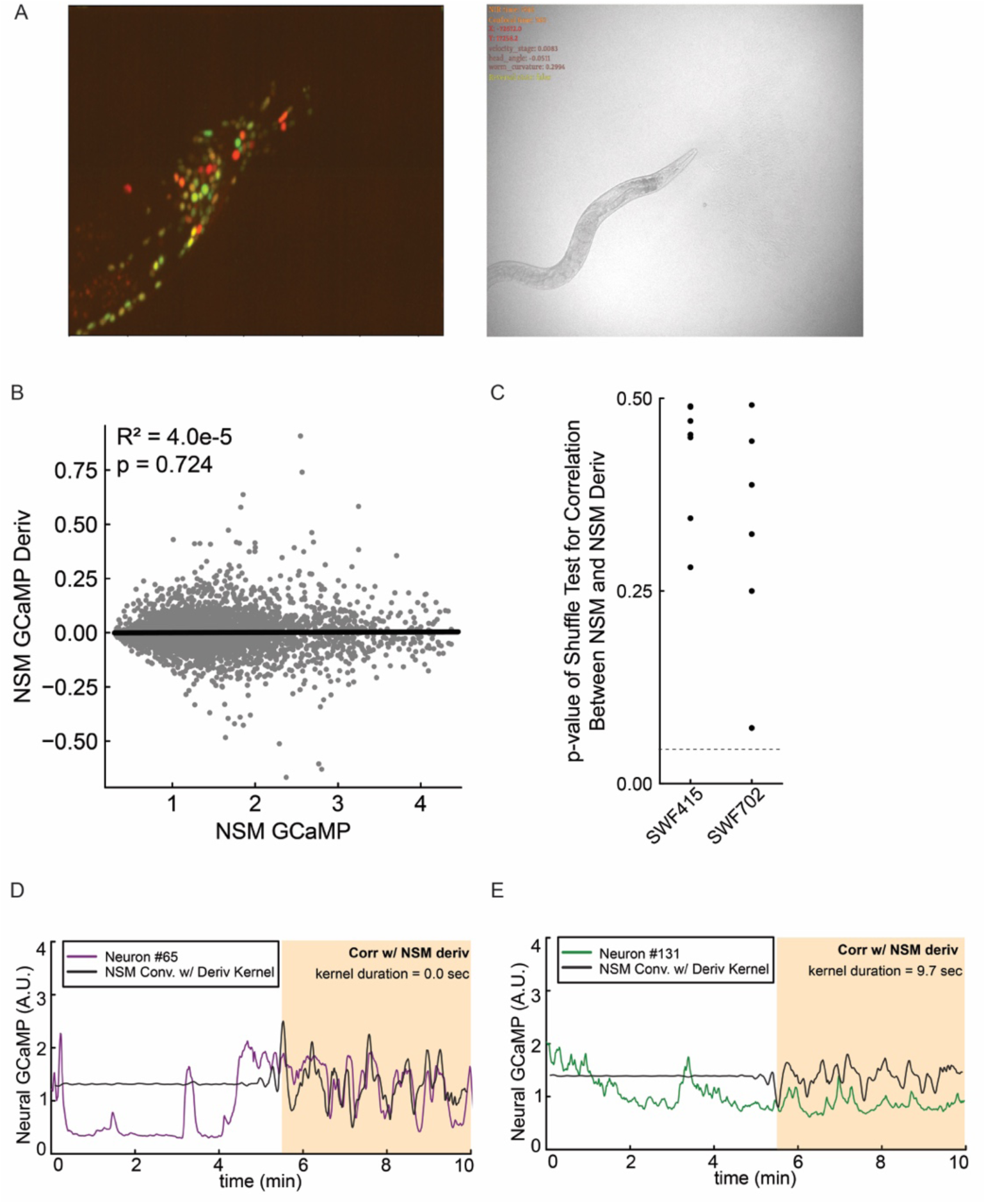
(A) Example images from a brain-wide recording. Left: A maximum intensity project of a single timepoint, showing the NLS-GCaMP7f (green) and NLS-mNeptune2.5 (red) signals in the head. Right: Behavioral image, captured in near infrared brightfield. (B) Scatterplot showing NSM activity and its derivative. Each dot is a timepoint and multiple animals are pooled together here. There is no significant correlation between the two. (C) Results of a statistical test to examine whether NSM’s activity was correlated with NSM derivative. This is related to panel B, except here we are performing the exact statistical test used throughout Figures 7 and 8 to test whether NSM itself is significantly correlated with its derivative (i.e. NSM activity convolved with filters that take its derivative). For all the animals recorded in Figure 7 (SWF415) and in Figure 8 (SWF702), there was no correlation. (D) Two example neurons that are also displayed in Fig. 7G that showed a significant positive (left) or negative (right) correlation with NSM’s derivative. Here, we show these neural traces again, but they are overlaid with NSM that has been convolved with a differentiator kernel (i.e. showing NSM derivative) so that this relationship can be more easily inspected. Note that the differentiator kernel on the right has a flipped sign, so that the black trace is basically the inverse of the derivative of NSM.

**Figure S8, Related to Figure 8.**
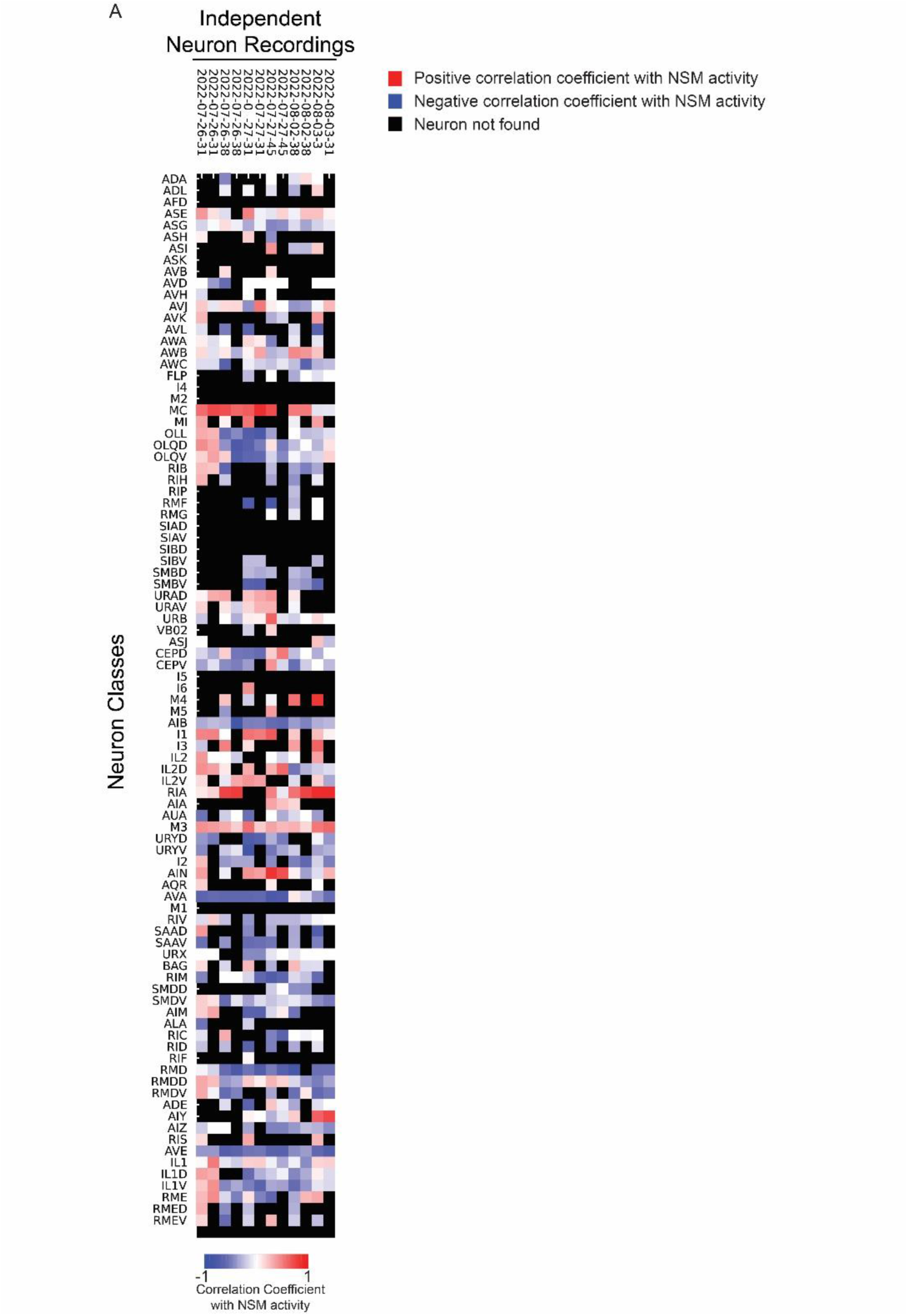
(A) Heat map depicting how each neuron class’ activity is associated with NSM activity in each recording. Neuron classes are shown as rows and the columns are the different recordings of that neuron (left and right neurons are shown separately). Pixels are black if the neuron was not found (or if the neuron class exists as a single cell).

## REFERENCES

Atanas, A.A., Kim, J., Wang, Z., Bueno, E., Becker, M., Kang, D., Park, J., Estrem, C., Kramer, T.S., Baskoylu, S., Mansingkha, V., Flavell, S.W., 2022. Brain-wide representations of behavior spanning multiple timescales and states in C. elegans. https://doi.org/10.1101/2022.11.11.516186

Béïque, J.-C., Imad, M., Mladenovic, L., Gingrich, J.A., Andrade, R., 2007. Mechanism of the 5-hydroxytryptamine 2A receptor-mediated facilitation of synaptic activity in prefrontal cortex. Proc. Natl. Acad. Sci. U. S. A. 104, 9870–9875. https://doi.org/10.1073/pnas.0700436104

Bocchio, M., McHugh, S.B., Bannerman, D.M., Sharp, T., Capogna, M., 2016. Serotonin, Amygdala and Fear: Assembling the Puzzle. Front. Neural Circuits 10, 24. https://doi.org/10.3389/fncir.2016.00024

Brittin, C.A., Cook, S.J., Hall, D.H., Emmons, S.W., Cohen, N., 2021. A multi-scale brain map derived from whole-brain volumetric reconstructions. Nature 591, 105–110. https://doi.org/10.1038/s41586-021-03284-x

Cazettes, F., Reato, D., Morais, J.P., Renart, A., Mainen, Z.F., 2021. Phasic Activation of Dorsal Raphe Serotonergic Neurons Increases Pupil Size. Curr. Biol. CB 31, 192–197.e4. https://doi.org/10.1016/j.cub.2020.09.090

Chase, D.L., Koelle, M.R., 2007. Biogenic amine neurotransmitters in C. elegans. WormBook Online Rev. C Elegans Biol. 1–15. https://doi.org/10.1895/wormbook.1.132.1

Cohen, J.Y., Amoroso, M.W., Uchida, N., 2015. Serotonergic neurons signal reward and punishment on multiple timescales. eLife 4. https://doi.org/10.7554/eLife.06346

Cook, S.J., Jarrell, T.A., Brittin, C.A., Wang, Y., Bloniarz, A.E., Yakovlev, M.A., Nguyen, K.C.Q., Tang, L.T.-H., Bayer, E.A., Duerr, J.S., Bülow, H.E., Hobert, O., Hall, D.H., Emmons, S.W., 2019. Whole-animal connectomes of both Caenorhabditis elegans sexes. Nature 571, 63–71. https://doi.org/10.1038/s41586-019-1352-7

Donaldson, Z.R., Nautiyal, K.M., Ahmari, S.E., Hen, R., 2013. Genetic approaches for understanding the role of serotonin receptors in mood and behavior. Curr. Opin. Neurobiol. 23, 399–406. https://doi.org/10.1016/j.conb.2013.01.011

Flavell, S.W., Pokala, N., Macosko, E.Z., Albrecht, D.R., Larsch, J., Bargmann, C.I., 2013. Serotonin and the neuropeptide PDF initiate and extend opposing behavioral states in C. elegans. Cell 154, 1023–1035. https://doi.org/10.1016/j.cell.2013.08.001

Goodfellow, N.M., Benekareddy, M., Vaidya, V.A., Lambe, E.K., 2009. Layer II/III of the prefrontal cortex: Inhibition by the serotonin 5-HT1A receptor in development and stress. J. Neurosci. Off. J. Soc. Neurosci. 29, 10094–10103. https://doi.org/10.1523/JNEUROSCI.1960-09.2009

Grossman, C.D., Bari, B.A., Cohen, J.Y., 2022. Serotonin neurons modulate learning rate through uncertainty. Curr. Biol. CB 32, 586–599.e7. https://doi.org/10.1016/j.cub.2021.12.006

Hobson, R.J., Hapiak, V.M., Xiao, H., Buehrer, K.L., Komuniecki, P.R., Komuniecki, R.W., 2006. SER-7, a Caenorhabditis elegans 5-HT7-like receptor, is essential for the 5-HT stimulation of pharyngeal pumping and egg laying. Genetics 172, 159–169. https://doi.org/10.1534/genetics.105.044495

Horvitz, H.R., Chalfie, M., Trent, C., Sulston, J.E., Evans, P.D., 1982. Serotonin and octopamine in the nematode Caenorhabditis elegans. Science 216, 1012–1014. https://doi.org/10.1126/science.6805073

Iwanir, S., Brown, A.S., Nagy, S., Najjar, D., Kazakov, A., Lee, K.S., Zaslaver, A., Levine, E., Biron, D., 2016. Serotonin promotes exploitation in complex environments by accelerating decision-making. BMC Biol. 14, 9. https://doi.org/10.1186/s12915-016-0232-y

Kennedy, A., Kunwar, P.S., Li, L.-Y., Stagkourakis, S., Wagenaar, D.A., Anderson, D.J., 2020. Stimulus-specific hypothalamic encoding of a persistent defensive state. Nature 586, 730–734. https://doi.org/10.1038/s41586-020-2728-4

Li, Y., Zhong, W., Wang, D., Feng, Q., Liu, Z., Zhou, J., Jia, C., Hu, F., Zeng, J., Guo, Q., Fu, L., Luo, M., 2016. Serotonin neurons in the dorsal raphe nucleus encode reward signals. Nat. Commun. 7, 10503. https://doi.org/10.1038/ncomms10503

Lottem, E., Banerjee, D., Vertechi, P., Sarra, D., Lohuis, M.O., Mainen, Z.F., 2018. Activation of serotonin neurons promotes active persistence in a probabilistic foraging task. Nat. Commun. 9, 1000. https://doi.org/10.1038/s41467-018-03438-y

Marques, J.C., Li, M., Schaak, D., Robson, D.N., Li, J.M., 2020. Internal state dynamics shape brainwide activity and foraging behaviour. Nature 577, 239–243. https://doi.org/10.1038/s41586-019-1858-z

Mathis, A., Mamidanna, P., Cury, K.M., Abe, T., Murthy, V.N., Mathis, M.W., Bethge, M., 2018. DeepLabCut: markerless pose estimation of user-defined body parts with deep learning. Nat. Neurosci. 21, 1281–1289. https://doi.org/10.1038/s41593-018-0209-y

McLachlan, I.G., Kramer, T.S., Dua, M., DiLoreto, E.M., Gomes, M.A., Dag, U., Srinivasan, J., Flavell, S.W., 2022. Diverse states and stimuli tune olfactory receptor expression levels to modulate food-seeking behavior. eLife 11, e79557. https://doi.org/10.7554/eLife.79557

Morud, J., Hardege, I., Liu, H., Wu, T., Choi, M.-K., Basu, S., Zhang, Y., Schafer, W.R., 2021. Deorphanization of novel biogenic amine-gated ion channels identifies a new serotonin receptor for learning. Curr. Biol. CB 31, 4282–4292.e6. https://doi.org/10.1016/j.cub.2021.07.036

Moyle, M.W., Barnes, K.M., Kuchroo, M., Gonopolskiy, A., Duncan, L.H., Sengupta, T., Shao, L., Guo, M., Santella, A., Christensen, R., Kumar, A., Wu, Y., Moon, K.R., Wolf, G., Krishnaswamy, S., Bao, Z., Shroff, H., Mohler, W.A., Colón-Ramos, D.A., 2021. Structural and developmental principles of neuropil assembly in C. elegans. Nature 591, 99–104. https://doi.org/10.1038/s41586-020-03169-5

Nelson, J.C., Colón-Ramos, D.A., 2013. Serotonergic neurosecretory synapse targeting is controlled by netrin-releasing guidepost neurons in Caenorhabditis elegans. J. Neurosci. Off. J. Soc. Neurosci. 33, 1366–1376. https://doi.org/10.1523/JNEUROSCI.3471-12.2012

Okaty, B.W., Commons, K.G., Dymecki, S.M., 2019. Embracing diversity in the 5-HT neuronal system. Nat. Rev. Neurosci. 20, 397–424. https://doi.org/10.1038/s41583-019-0151-3

Okaty, B.W., Sturrock, N., Escobedo Lozoya, Y., Chang, Y., Senft, R.A., Lyon, K.A., Alekseyenko, O.V., Dymecki, S.M., 2020. A single-cell transcriptomic and anatomic atlas of mouse dorsal raphe Pet1 neurons. eLife 9, e55523. https://doi.org/10.7554/eLife.55523

Olde, B., McCombie, W.R., 1997. Molecular cloning and functional expression of a serotonin receptor from Caenorhabditis elegans. J. Mol. Neurosci. MN 8, 53–62. https://doi.org/10.1007/BF02736863

Paquelet, G.E., Carrion, K., Lacefield, C.O., Zhou, P., Hen, R., Miller, B.R., 2022. Single-cell activity and network properties of dorsal raphe nucleus serotonin neurons during emotionally salient behaviors. Neuron 110, 2664–2679.e8. https://doi.org/10.1016/j.neuron.2022.05.015

Puig, M.V., Gulledge, A.T., 2011. Serotonin and prefrontal cortex function: neurons, networks, and circuits. Mol. Neurobiol. 44, 449–464. https://doi.org/10.1007/s12035-011-8214-0

Puig, M.V., Watakabe, A., Ushimaru, M., Yamamori, T., Kawaguchi, Y., 2010. Serotonin modulates fast-spiking interneuron and synchronous activity in the rat prefrontal cortex through 5-HT1A and 5-HT2A receptors. J. Neurosci. Off. J. Soc. Neurosci. 30, 2211– 2222. https://doi.org/10.1523/JNEUROSCI.3335-09.2010

Ranganathan, R., Cannon, S.C., Horvitz, H.R., 2000. MOD-1 is a serotonin-gated chloride channel that modulates locomotory behaviour in C. elegans. Nature 408, 470–475. https://doi.org/10.1038/35044083

Ren, J., Friedmann, D., Xiong, J., Liu, C.D., Ferguson, B.R., Weerakkody, T., DeLoach, K.E., Ran, C., Pun, A., Sun, Y., Weissbourd, B., Neve, R.L., Huguenard, J., Horowitz, M.A., Luo, L., 2018. Anatomically Defined and Functionally Distinct Dorsal Raphe Serotonin Sub-systems. Cell 175, 472–487.e20. https://doi.org/10.1016/j.cell.2018.07.043

Ren, J., Isakova, A., Friedmann, D., Zeng, J., Grutzner, S.M., Pun, A., Zhao, G.Q., Kolluru, S.S., Wang, R., Lin, R., Li, P., Li, A., Raymond, J.L., Luo, Q., Luo, M., Quake, S.R., Luo, L., 2019. Single-cell transcriptomes and whole-brain projections of serotonin neurons in the mouse dorsal and median raphe nuclei. eLife 8, e49424. https://doi.org/10.7554/eLife.49424

Rhoades, J.L., Nelson, J.C., Nwabudike, I., Yu, S.K., McLachlan, I.G., Madan, G.K., Abebe, E., Powers, J.R., Colón-Ramos, D.A., Flavell, S.W., 2019. ASICs Mediate Food Responses in an Enteric Serotonergic Neuron that Controls Foraging Behaviors. Cell 176, 85–97.e14. https://doi.org/10.1016/j.cell.2018.11.023

Sawin, E.R., Ranganathan, R., Horvitz, H.R., 2000. C. elegans locomotory rate is modulated by the environment through a dopaminergic pathway and by experience through a serotonergic pathway. Neuron 26, 619–631. https://doi.org/10.1016/s0896-6273(00)81199-x

Seo, C., Guru, A., Jin, M., Ito, B., Sleezer, B.J., Ho, Y.-Y., Wang, E., Boada, C., Krupa, N.A., Kullakanda, D.S., Shen, C.X., Warden, M.R., 2019. Intense threat switches dorsal raphe serotonin neurons to a paradoxical operational mode. Science 363, 538–542. https://doi.org/10.1126/science.aau8722

Taylor, S.R., Santpere, G., Weinreb, A., Barrett, A., Reilly, M.B., Xu, C., Varol, E., Oikonomou, P., Glenwinkel, L., McWhirter, R., Poff, A., Basavaraju, M., Rafi, I., Yemini, E., Cook, S.J., Abrams, A., Vidal, B., Cros, C., Tavazoie, S., Sestan, N., Hammarlund, M., Hobert, O., Miller, D.M., 2021. Molecular topography of an entire nervous system. Cell 184, 4329–4347.e23. https://doi.org/10.1016/j.cell.2021.06.023

Towlson, E.K., Vértes, P.E., Ahnert, S.E., Schafer, W.R., Bullmore, E.T., 2013. The rich club of the C. elegans neuronal connectome. J. Neurosci. Off. J. Soc. Neurosci. 33, 6380–6387. https://doi.org/10.1523/JNEUROSCI.3784-12.2013

Weissbourd, B., Ren, J., DeLoach, K.E., Guenthner, C.J., Miyamichi, K., Luo, L., 2014. Presynaptic partners of dorsal raphe serotonergic and GABAergic neurons. Neuron 83, 645–662. https://doi.org/10.1016/j.neuron.2014.06.024

White, J.G., Southgate, E., Thomson, J.N., Brenner, S., 1986. The structure of the nervous system of the nematode Caenorhabditis elegans. Philos. Trans. R. Soc. Lond. B. Biol. Sci. 314, 1–340. https://doi.org/10.1098/rstb.1986.0056

Witvliet, D., Mulcahy, B., Mitchell, J.K., Meirovitch, Y., Berger, D.R., Wu, Y., Liu, Y., Koh, W.X., Parvathala, R., Holmyard, D., Schalek, R.L., Shavit, N., Chisholm, A.D., Lichtman, J.W., Samuel, A.D.T., Zhen, M., 2021. Connectomes across development reveal principles of brain maturation. Nature 596, 257–261. https://doi.org/10.1038/s41586-021-03778-8

Yemini, E., Lin, A., Nejatbakhsh, A., Varol, E., Sun, R., Mena, G.E., Samuel, A.D.T., Paninski, L., Venkatachalam, V., Hobert, O., 2021. NeuroPAL: A Multicolor Atlas for Whole-Brain Neuronal Identification in C. elegans. Cell 184, 272–288.e11. https://doi.org/10.1016/j.cell.2020.12.012

Yohn, C.N., Gergues, M.M., Samuels, B.A., 2017. The role of 5-HT receptors in depression. Mol. Brain 10, 28. https://doi.org/10.1186/s13041-017-0306-y

